# A robust deep neural network for denoising task-based fMRI data: An application to working memory and episodic memory

**DOI:** 10.1101/746313

**Authors:** Zhengshi Yang, Xiaowei Zhuang, Karthik Sreenivasan, Virendra Mishra, Tim Curran, Dietmar Cordes

**Author notes:** Correspondence to: Dietmar Cordes, Ph.D., Cleveland Clinic Lou Ruvo Center for Brain Health 888 W. Bonneville Ave, Las Vegas, NV, 89106.

## Abstract

In this study, a deep neural network (*DNN*) is proposed to reduce the noise in task-based fMRI data without explicitly modeling noise. The *DNN* artificial neural network consists of one temporal convolutional layer, one long short-term memory (LSTM) layer, one time-distributed fully-connected layer, and one unconventional *selection* layer in sequential order. The LSTM layer takes not only the current time point but also what was perceived in a previous time point as its input to characterize the temporal autocorrelation of fMRI data. The fully-connected layer weights the output of the LSTM layer, and the output denoised fMRI time series is selected by the *selection* layer. Assuming that task-related neural response is limited to gray matter, the model parameters in the *DNN* network are optimized by maximizing the correlation difference between gray matter voxels and white matter or ventricular cerebrospinal fluid voxels. Instead of targeting a particular noise source, the proposed neural network takes advantage of the task design matrix to better extract task-related signal in fMRI data. The *DNN* network, along with other traditional denoising techniques, has been applied on simulated data, working memory task fMRI data acquired from a cohort of healthy subjects and episodic memory task fMRI data acquired from a small set of healthy elderly subjects. Qualitative and quantitative measurements were used to evaluate the performance of different denoising techniques. In the simulation, DNN improves fMRI activation detection and also adapts to varying hemodynamic response functions across different brain regions. DNN efficiently reduces physiological noise and generates more homogeneous task-response correlation maps in real data.

## 1. INTRODUCTION

Functional magnetic resonance imaging (fMRI) based on the blood-oxygen-level dependent (Kasper et al.) signal has been widely used to investigate brain function in human research. The BOLD signal is an indirect measure of neuronal activity and is contaminated by a large proportion of noise fluctuations (Bianciardi et al., 2009; Caballero-Gaudes and Reynolds, 2017). Noise fluctuations arise from multiple sources such as thermal noise inherent to electrical circuits, head-motion-related signal change, cardiac and respiratory oscillations, instrumental drift, changes in blood pressure, and cerebral autoregulation mechanisms (Murphy et al., 2013). These noise sources can considerably affect the result and interpretation of any task-based and resting-state fMRI experiments. Separating signal from noise in fMRI data leads to improved signal-to-noise ratio (SNR) and statistical power of fMRI data analysis. In this study, we focus on denoising task-based fMRI data to better identify task-involved brain regions.

Nuisance regression has been extensively used to capture non-neural fluctuation and reduce noise in fMRI time series (Friston et al., 1995). One way to apply nuisance regression is to first extract nuisance regressors from fMRI data or externally recorded data, and then apply the general linear model (GLM) to obtain denoised data. Alternatively, specify the nuisance regressors in one design matrix as additional regressors to account for their variance in fMRI activation analysis. The degrees of freedom, when nuisance regression is applied, needs to be properly adjusted to account for the additional regressors in the statistical analysis. Denoising methods based on nuisance regression have been mainly developed aiming at correcting the effect of head motion or major physiological noise fluctuations, such as cardiac and respiratory contaminations.

Motion correction methods usually treat six motion parameters (R=[X Y Z pitch yaw roll]) estimated from rigid-body affine transformation as nuisance regressors to address motion-related noise in fMRI data (Friston et al., 1996; Johnstone et al., 2006). Extensions of this method were also developed by including the derivatives (R’) and squares (R^2^) of motion parameters to remove spin history related aspects of motion-related artifacts (Friston et al., 1996). Motion parameters and their variants, however, may not be sufficient to model motion-related artifacts in fMRI data. Moreover, increasing the number of nuisance regressors can also substantially remove the BOLD signal.

Cardiac and respiratory noise fluctuations can explain considerable amount of the variance in task-based fMRI data (Bianciardi et al., 2009; Jorge et al., 2013; Triantafyllou et al., 2011). One study showed that physiological fluctuations could explain about 10-15% variance of their fMRI data (Cordes et al., 2014). One way to reduce these noise sources is by externally recording the spectrum of cardiac and respiratory oscillations, identifying the fundamental frequency of these fluctuations, and applying notch filters at the fundamental frequencies. The direct side effect of filtering is that any BOLD signal at these frequencies is also reduced by the filter (Biswal et al., 1996). Consequently, more sophisticated modeling approaches (Birn et al., 2008; Chang et al., 2009; Glover et al., 2000; Harvey et al., 2008; Shmueli et al., 2007; Tijssen et al., 2014) were later proposed to reduce the main effects of cardiac and respiratory fluctuations with externally recorded physiological wave functions.

Along with the denoising methods explicitly modeling a particular noise source, data-driven methods were also developed in the last decade without assuming any parametric noise model or externally recorded wave functions. Using the average signals of white matter (WM) and cerebrospinal fluid (CSF) for nuisance regression is a popular and simple data-driven method (Anderson et al., 2011; Jo et al., 2010). Behzadi et al. (2007) proposed a principal component analysis (PCA) based method, CompCor, which derives nuisance regressors from the regions where no neuronal signal is expected. CompCor was shown to be an efficient method to reduce physiological noise without external recordings. Another PCA-based denoising method GLMdenoise (Kay et al., 2013) applies PCA on task-unrelated voxels after an initial fitting and then uses cross-validation to estimate the optimal number of principal components as nuisance regressors. The cross-validation step in GLMdenoise requires multiple runs for the same task on a single subject, which are usually not available in most datasets. Another strategy to denoise fMRI data is based on independent component analysis. The independent components (ICs) first can be classified as noise or signal components by manual labelling, and then the noise components are regressed out from fMRI data. Since manual labelling is time-consuming and subjective, the classifiers such as FIX (Salimi-Khorshidi et al., 2014) and ICA-AROMA (Pruim et al., 2015) were introduced to distinguish ICs with less human intervention. These methods can be used as automated techniques if a new dataset is denoised by a classifier pre-trained with manually labeled ICs. If a particular dataset is substantially different from pre-trained data, the classifier should be trained by researchers themselves after manually labelling the ICs from a few subjects. More detailed discussion about the methods mentioned above can be seen in the denoising review paper (Caballero-Gaudes and Reynolds, 2017). In addition to the *post-hoc* methods mentioned above, multi-echo EPI sequence was recently proposed to differentiate BOLD and non-BOLD signal during data acquisition by investigating the echo time dependence of the ICA components (Kundu et al., 2012). Only BOLD components have a linear dependence as a function of the echo time.

In this study, we proposed a robust and automated deep neural network, *DNN*, for denoising task-based fMRI data. *DNN* is a data-driven method that does not assume a specific noise model and optimizes its parameters for each individual subject without expert intervention. The *DNN* network consists of a convolutional layer, a long short-term memory (LSTM) layer (Hochreiter and Schmidhuber, 1997), a time-distributed fully-connected layer, and a nonconventional *selection* layer. The convolutional layer can be treated as an adaptive temporal filter. Artificial neural networks constructed with LSTM layers have achieved accuracy records in analyzing time-sequential data such as language modeling, speech recognition, and machine translation (Bahdanau et al., 2014; Graves et al., 2013; Sundermeyer et al., 2012). The LSTM in this study is a neural network that takes not only the current time point but also the previous time point as input to account for the temporal autocorrelation in fMRI time-series data. A time-distributed fully-connected layer is used to optimize the weights of the multiple outputs from the LSTM layer, and the *selection* layer determines the output time series for each voxel. More detail on the architecture of the denoising network is provided in the *Theory* section.

The *DNN* network, along with other denoising techniques, has been applied on simulated data, working memory task fMRI data acquired from a cohort of healthy subjects and episodic memory task fMRI data acquired from a small set of healthy elderly subjects.

## 2. THEORY

### 2.1. Architecture of the *DNN* network

The *DNN* network consists of four layers in a sequential order, namely a 1-dimensional (1-dim) convolutional layer, a LSTM layer, a time-distributed fully-connected layer and a *selection* layer. Each voxel is treated as a sample and each time point is treated as a feature. Let us denote *N* as the number of voxels and *T* as the length of the time series, then the input to the network are the minimum-preprocessed fMRI data with dimension of *N* x *T* x 1, where the singleton dimension “1” represents the number of channels following the terminology in the deep learning community.

The purpose of the *DNN* network is to reduce noise in the fMRI data with multiple layers and then output the denoised data. Naturally, the output layer has the same dimension as the input layer. If *V* 1-dim filters are used in the convolutional layer, *L* units are used in the LSTM layer and *K* nodes are specified in time-distributed fully-connected layer. The output dimensions from these layers are *N* x *T* x *V*, *N* x *T* x *L* and *N* x *T* x *K*, respectively. There are *K* time series for a single voxel after the time-distributed fully-connected layer. The *selection* layer is a nonconventional layer without any parameters and its purpose is to choose a single time series from the *K* time series for each voxel having maximal correlation with the task design matrix.

### 2.2. 1-dim convolutional layer

Since the input features to the convolutional layer are the time points of fMRI data, the 1-dim convolutional layer can be treated as a set of temporal filters. In standard fMRI preprocessing steps, a high-pass filter is commonly used to remove slow-varying signal drifts less than 0.01Hz from fMRI time series data and a low-pass filter is used to capture the low-frequency BOLD signal less than 0.1Hz under the hypothesis that there is no significant high-frequency BOLD signal above 0.1Hz (Cordes et al., 2001). However, the frequency threshold is still debatable and a few recent studies point out that BOLD fractional contribution may extend above 0.1 Hz (Boubela et al., 2013; Chen and Glover, 2015). In addition, the frequency threshold may be subject-specific. In contrast, there is no frequency threshold specified in the convolutional layer of the *DNN* and the temporal filters are learned from the data itself without expert’s intervention. Since the *DNN* is carried out for each subject independently, the convolutional layer estimates a set of subject-specific temporal filters for data denoising. The parameters that needs to be optimized in the 1-dimensional convolutional layer is a set of filters ***W***_conv_ ∈ *R*^S×V^, where *S* represents the filter size and *V* represents the number of filters, as explained in section 2.1. Fig.2a shows how a 1-dim filter is applied on a time series with filter size 5 and stride length 1. The filter is applied on the time points limited by the filter size (green box) to output a single time point and then shifts one time point (red box) to generate the next time point. The same process is repeated *T* times to output one time series. Naturally, the convolutional layer ***W***_conv_ with *V* filters applied on a single time series has output time series with size of *T* x *V*.

**Figure 1.**
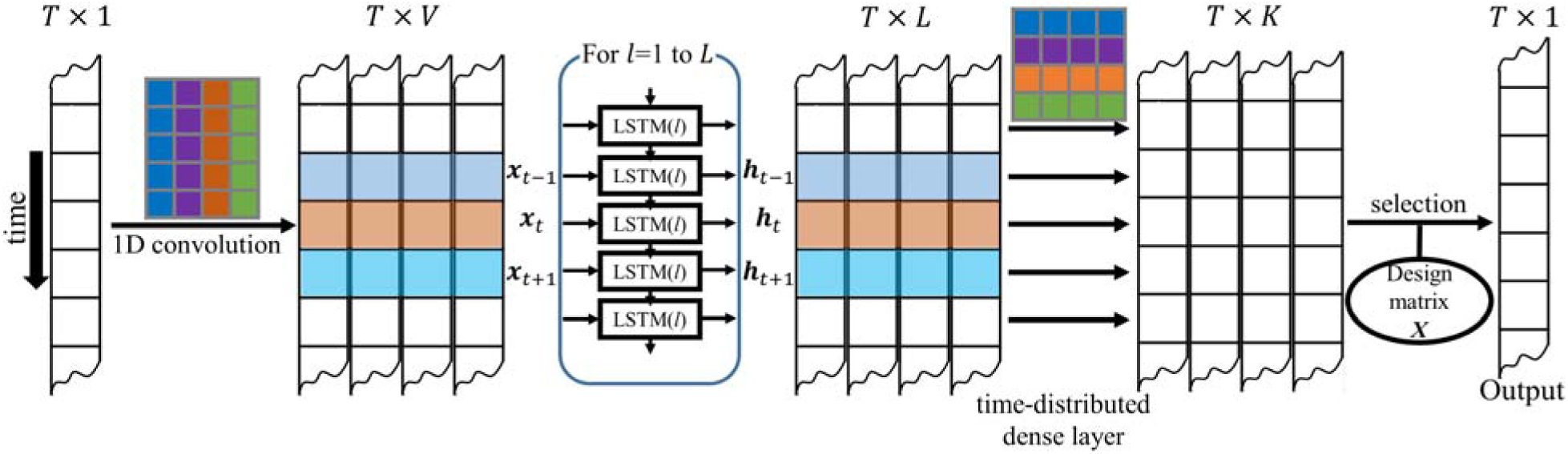
The architecture of the *DNN* network. In sequential order, the *DNN* network consists of a 1-dimensional convolutional layer (with respect to time) with *V* temporal filters (*V*=4, filter size f=5, stride s=1), an LSTM layer with *L* units (*L*=4), and a time-distributed fully-connected layer with *K* nodes (*K*=4) and a *selection* layer. There are *K* time series for a single voxel after the time-distributed fully-connected layer. The selection layer is a nonconventional layer without any parameters and its purpose is to choose a single time series from the *K* time series for each voxel having maximal correlation with the task design matrix *X*.

**Figure 2.**
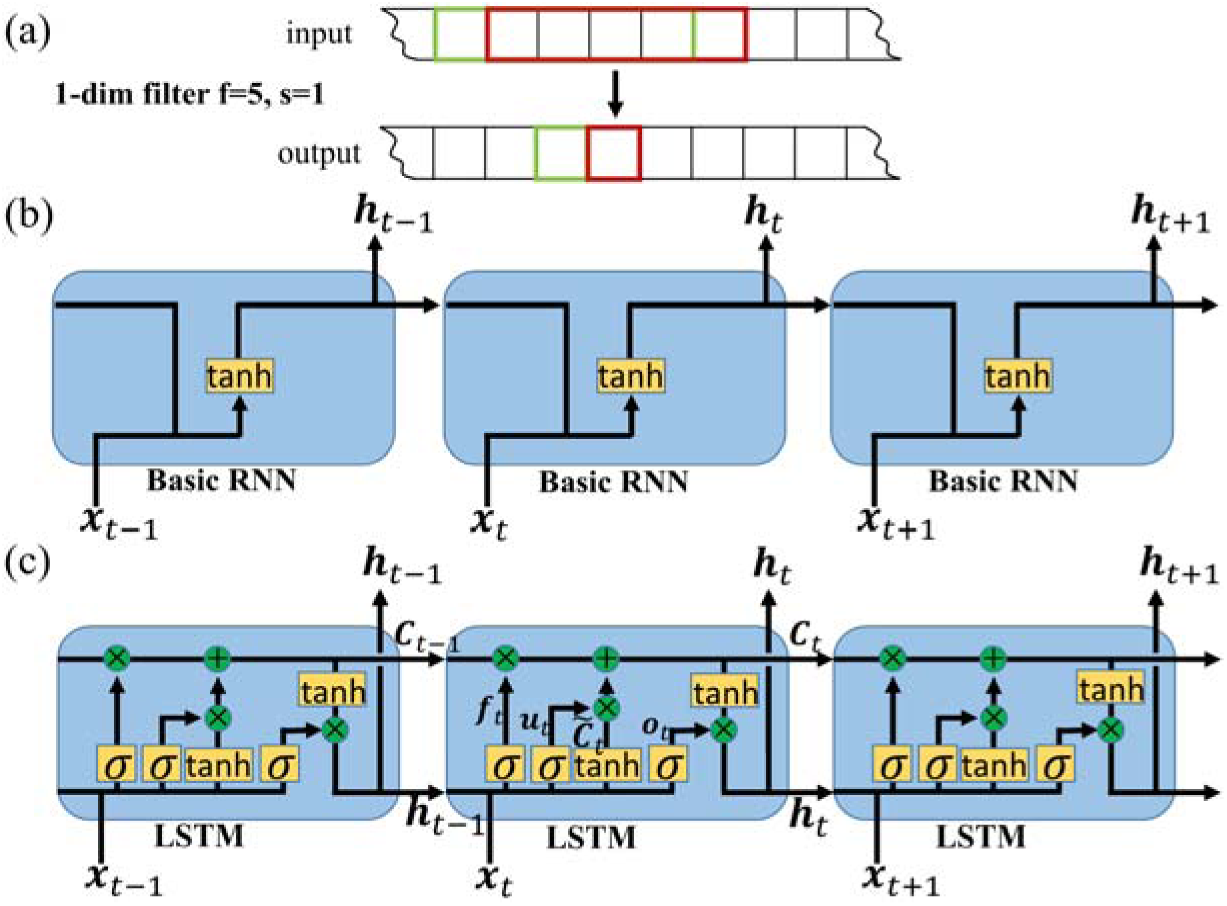
(a) Example of a filter in 1-dim convolutional layer with filter size 5 and stride length 1. Subfigures (b) and (c) show a comparison of basic RNN layer (b) and LSTM layer (c) with a hyperbolic tangent as an activation function.

### 2.3. LSTM layer

4D fMRI data represent a time sequence of 3D brain imaging data. Due to temporal autocorrelations of the BOLD signal, the fMRI response at a current time point is partially depended to the previous time point. Similar to a human who interprets each word based on previous words in a sentence, a LSTM network is one special kind of recurrent neural networks (RNNs) that takes the information from previous time points to inform the current time point.

Specifically, RNN networks including LSTM take the temporal autocorrelation of the data into consideration during the training of the networks. The chain-like structure in Fig.2b shows that RNN networks are closely related to sequences or lists, such as time series in fMRI data. Because a basic RNN network as in Fig.2b has difficulty in handling long-term dependencies (Bengio et al., 1994), LSTM networks were introduced by Hochreiter and Schmidhuber (1997) and are now widely used in the deep learning community for analyzing time-serial data. The basic RNN network has a simple structure consisting of an activation function applied to the output from the previous time point ***h***_t-1_ and the current time point ***x***_t_. The schematic diagram of the LSTM layer is shown in Fig.2c. The horizontal line running through the top of the diagram connects all cell states ***C***_*t*_, *t* = 1,…, *T*. The cell states run straight across the entire sequence, and thus it is easy for previous information to flow to the next time points. The LSTM is able to add or remove information to cell states that are carefully controlled by three gates called *forget gate*, *update gate* and *output gate*. These gates consist of a sigmoid neural net layer represented as 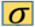 and a pointwise multiplication operation represented as 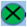 in Fig.2c. The sigmoid layers have output values ranging from zero to one, which describe the percentage of information that should be let through. Following the notation used in the deep learning community, the coefficient matrices ***f***_t_, ***u***_t_, and ***o***_t_ are defining the output from the sigmoid layers from *forget gate*, *update gate*, and *output gate*, respectively. The coefficient matrices (vectors for each time point *t*) depend on weight matrices ***W***_f_, ***W***_u_, and ***W***_o_ and bias vectors ***b***_f_, ***b***_u_, and ***b***_o_, namely,

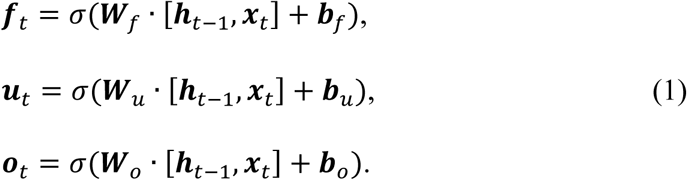

The bias vectors in Eq. (1) are broadcasted (in python terminology) into matrices to match the data dimension during computation. The matrix ***f***_t_ (with same dimension as ***C***_t-1_) contains a value between zero and one for each number in the cell state ***C***_t-1_. A value of zero in ***f***_t_ means “completely remove this number”. The pointwise multiplication ***f***_t_ X ***C***_t-1_ represents how much information from the previous cell state ***C***_t-1_ remains for the new cell state ***C***_t_. The coefficient matrix ***u***_t_ computed from the second sigmoid layer scales the output ***C*** _t_ from hyperbolic tangent layer with a coefficient matrix ***W***_c_ and then a pointwise addition is used to update the new cell state ***C***_t_,

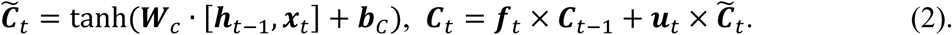

In addition to the information from the previous cell state, the new cell state ***C***_t_ is updated with the current input ***x***_t_. Finally, the output ***h***_t_ in LSTM is calculated based on the cell state ***C***_t_ with the hyperbolic tangent function as activation function and scaled with ***o***_t_ from the output gate,

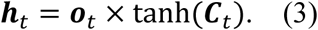

### 2.4. Time-distributed fully-connected layer

The third layer in the *DNN* model is the time-distributed fully-connected layer with an input of dimension *N* x *T* x *L* from the LSTM layer. The standard fully-connected layer connects all input nodes with the output nodes ignoring the sequential property of the data (left panel of Fig.3). The weight matrix has a dimension of *K* x (*T* x *L*) and hence the output from the fully-connected layer has the dimension *N* x *K* if *K* nodes are specified. In contrast, a time-distributed fully-connected layer applies the same fully-connected operation to every time point in the data. The weight matrix in the fully-connected layer has the dimension *K* x *L*, and the output has the dimension *N* x *T* x *K* which is the same as the input fMRI data.

**Figure 3.**
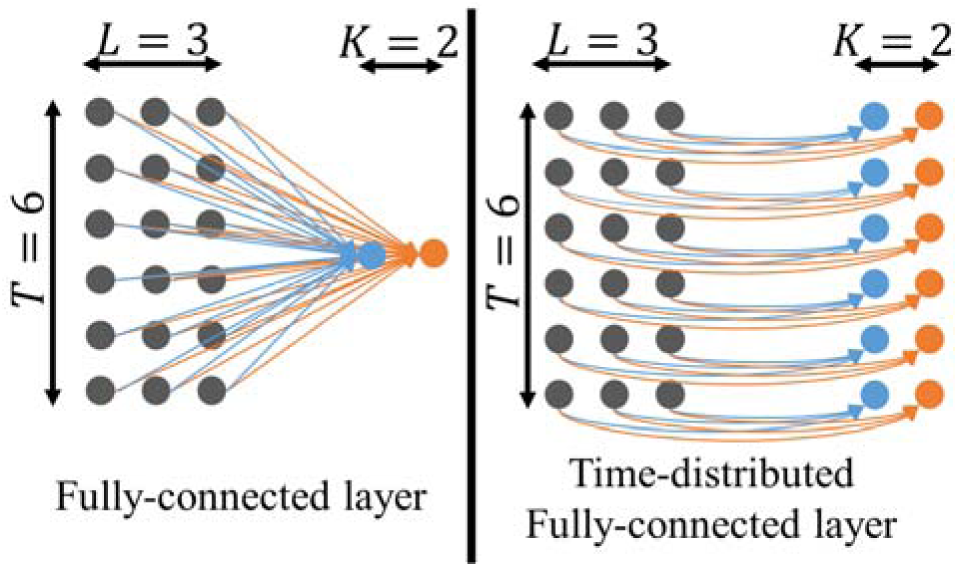
Schematic diagrams of fully-connected layer (left) and time-distributed fully-connected layer (right).

### 2.5. *Selection* layer

The *selection* layer is designed to select a single time series from the output of *K* time series of the time-distributed fully-connected layer. The correlation between each time series and the design matrix is calculated using the GLM (see Eq.4) and the one time series having maximal correlation is selected as the output for each voxel. In standard fMRI activation analysis, the task regressor in the design matrix is chosen by convolving the binary task with the canonical hemodynamic response function (HRF) without considering that the HRF may differ across brain regions. However, the time-distributed fully-connected layer with multiple nodes allows the neural network to adapt to different HRFs and the selection layer chooses a single time series in an optimal manner. If the time-distributed fully-connected layer is specified with a single node, namely *K* = 1, the output from the LSTM layer is directly the denoised time series and the selection layer can be skipped.

### 2.6. Customized cost function for *DNN* network

The cost function provides a criterion to update all parameters in the network during the iteration step. Many cost functions have been developed for the purpose of classification or regression in machine learning applications, such as mean squared error, mean absolute percentage error, cross entropy, Poisson, and cosine proximity cost functions. These cost functions are calculated with the known true values or classes, however, the challenge to construct a cost function for *DNN* denoising is that the true BOLD signal in fMRI data is not available. For this reason, we have proposed a customized cost function which does not require knowing the true BOLD signal.

The DNN network is trained by first arbitrarily pairing one gray matter voxel with one white matter or ventricle voxel and then assigning the paired voxels to different batches. In each batch, let ***Y***_raw_ denotes the original fMRI data within the gray matter mask (GM mask) and ***Ỹ*** _raw_ denotes the paired time series within eroded white matter or ventricle mask (nonGM mask). During the learning process, ***Y***_raw_ and ***Ỹ*** _raw_ share exactly the same *DNN* network and are input to the network alternatingly for each iteration. Each iteration will then provide the corresponding output data ***Y***_*denoise*_ and ***Ỹ***_*denoise*_. The cost function is calculated with three variables, namely, ℒ = ℒ(***Y***_denoise_, ***Y*** _denoise_, ***X***) where ***X*** is the task-related design matrix. The cost function is defined as the correlation coefficient (*r*) between denoised time series within GM (***y***) and nonGM (***ỹ***) masks with the task design matrix ***X*** given by

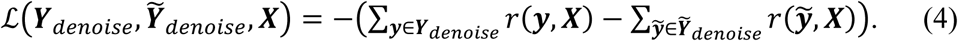

The correlation coefficient is calculated based on the GLM model ***y*** = ***Xβ*** + **ϵ** where ***β*** = ***X***^+^***y*** and **ϵ** is a Gaussian distributed random error term. The term *r*(***ỹ***, ***X***) is defined as the Pearson correlation between ***y*** and ***Xβ***. A similar definition is used for *r*(***ỹ***, ***X***). Because the squared correlation coefficient *r*(***ỹ***, ***X***)^2^ represents the proportion of the variance in the denoised time series ***y*** that can be explained by the task design matrix ***X***, the loss function can also be thought to be the difference of the square root of the task-explained variance in the denoised time series.

Since the voxels within nonGM masks (including eroded WM and CSF masks) are expected to have no neural response, the optimal network parameters are obtained by maximizing the correlation difference between voxels in GM and nonGM, leading to a minimum cost function. Considering that pseudoinverse ***X***^+^ of regressor ***X*** is required in solving the GLM and ***X***^+^ is independent of all other variables except ***X*** itself, ***X***^+^ is pre-calculated and added as a variable in the cost function, i.e. ℒ = ℒ(***Y***_*denoise*_, ***Ỹ***_*denoise*_, ***X***, ***X***^+^). In this case, there are no expensive inverse operations necessary when calculating the cost function, which makes the network more computationally efficient.

### 2.7. *DNN* architecture, model training and calibration

Since GM and nonGM voxels are paired to calculate the customized loss function, the same number of voxels within GM and nonGM masks are required to be the input samples to the network. During the optimization, the voxels within GM are randomly paired with voxels within nonGM and the extra voxels within GM or nonGM are discarded for the following optimization. 90% of the paired-voxels are randomly assigned to the training set to update parameters and the remaining 10% of the paired-voxels are assigned to the validation set to monitor whether the network suffers from over-fitting or under-fitting leading to high bias or variance, respectively. The initial parameters are randomly sampled from the Xavier uniform initializer (Glorot and Bengio, 2010). The parameters are updated with the *Adam* stochastic gradient-based optimization algorithm (Kingma and Ba, 2014), which adapts the parameter learning rates by taking advantage of both the average first moment (mean) and the average of the second moments of the gradients (uncentered variance). The *Adam* optimizer is parameterized with learning rate *η*=0.01, learning rate decay *γ*=0.05, exponential decay rate for the first moment estimates *β*_1_ = 0.9 and exponential decay rate for the second moments estimates *β*_2_ = 0.999. The parameters and their learning gradients are computed with each batch of 500 samples, and one epoch is defined as running through all batches.

## 3. MATERIALS and METHODS

### 3.1. Subjects

The structural and functional MRI data were obtained from the Human Connectome Project (HCP) database (https://www.humanconnectome.org/). The HCP project (Principal Investigators: Bruce Rosen, M.D., Ph.D., Martinos Center at Massachusetts General Hospital; Arthur W. Toga, Ph.D., University of Southern California, Van J. Weeden, MD, Martinos Center at Massachusetts General Hospital) is supported by the National Institute of Dental and Craniofacial Research (NIDCR), the National Institute of Mental Health (NIMH) and the National Institute of Neurological Disorders and Stroke (NINDS). HCP is the result of efforts of co-investigators from the University of Southern California, Martinos Center for Biomedical Imaging at Massachusetts General Hospital (MGH), Washington University, and the University of Minnesota.

The 3T MRI imaging data from 88 subjects were used in this study. All subjects are males with age in the range of 26-30 years having complete T1, resting-state MRI and task fMRI scans. The structural T1 images were acquired with a resolution of 260 x 311 x 260 to yield 0.7 mm x 0.7 mm x 0.7 mm isotropic voxel size. Resting-state fMRI data were acquired with 1,200 time frames from a gradient-echo fast EPI sequence with parameters: multiband factor 8, TR/TE=720/33.1 ms; flip angle=52 degrees; 72 slices; spatial resolution=2 mm x 2 mm x 2 mm and imaging matrix=104 x 90. There are seven tasks for each subject but we focus only on the working memory task fMRI study. The working memory task fMRI data were acquired with identical pulse sequence settings as resting-state fMRI but with 405 time points. The first 15 volumes of fMRI data were discarded to avoid data with unsaturated T1 signal. The minimally preprocessed resting-state fMRI data and the working memory task fMRI data (in standard MNI space) with additional linear detrending step were treated as raw fMRI data in our analysis. A more detailed description about HCP minimal preprocessing steps can be found in Glasser et al. (2013). The task itself represents an event-related task design consisting of targets, non-targets, and lures contrasts suggested by HCP site investigators was used in this study. Within each working memory task fMRI run, four different stimulus types including places, tools, faces, and body parts were presented in separate blocks. ½ of the blocks use a 2-back working memory task and ½ use a 0-back working memory task for each run. Further details of the HCP 3T MRI protocols and task designs can be found on the HCP website. Each subject’s T1 image was segmented into three tissue masks for GM, WM and CSF. To reduce partial volume effects, CSF was eroded once and WM was eroded up to four times (Power et al., 2014) with additional condition that the eroded WM mask has at least 10,000 voxels to ensure enough samples to train the neural network. The average volume of the eroded WM and CSF masks were 139 cm^3^ (17,457 voxels, 77% of WM voxels were removed by erosion) and 14 cm^3^ (1,726 voxels, 93% of CSF voxels were removed by erosion), respectively. The eroded masks were used to confine nonGM voxels used in DNN denoising and compute CompCor (Behzadi et al., 2007) regressors.

### 3.2. Simulation: Uniform HRF (uniHRF) and variable HRF (varHRF)

The simulation was generated to mimic real fMRI data for evaluating the performance of different denoising techniques. Considering that resting-state fMRI data share similar noise as in task fMRI data but without neural signal corresponding to a specific task, the basic idea behind the simulation is to add task-related signal to resting-state data. 15 task fMRI datasets were simulated by adding signal to resting-state data from 15 subjects. Since the resting-state data had a total of 1200 volumes but the working memory task fMRI data had only 390 volumes after discarding the first 15 volumes, the resting-state data were truncated and only the frames from 201 to 590 were used for the simulation. The binary working memory task design consisting of targets, non-targets and lures was also extracted for each subject. 6 bilateral AAL regions (Tzourio-Mazoyer et al., 2002), including anterior cingulate cortex, precentral gyrus, inferior frontal gyrus, insula, middle frontal gyrus and middle temporal gyrus, were specified to be active. Two sets of data were simulated. The signal in the first dataset was generated with the canonical HRF (uniHRF) and the signal in the second dataset was generated with varying HRFs for these six active regions (varHRF). Given the fact that HRFs are heterogeneous by having different onset delays across brain regions, the varHRF simulation is useful to evaluate the sensitivity of our proposed *DNN* method to potential misspecification of heterogeneous HRFs in the estimated task effects. The signal within each active region was generated by the following steps:

The weights in the six active regions can be arbitrarily defined as long as these three possible senarios are covered: 1) a region only involved in one condition (e.g. [1 0 0]^T^); 2) a region involved in two conditions (e.g. [0.3 1 0]^T^); 3) a region involved in all three conditions (e.g. [1 0.3 0.3]^T^). The set of weights *B* ∈ ℛ^3X6^ for our simulation was specified as

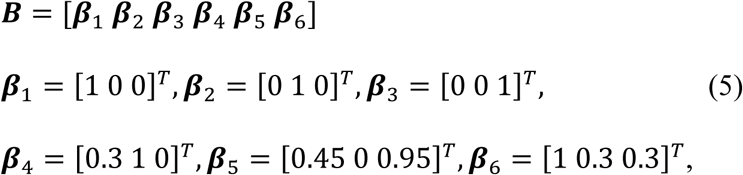

with the criteria described above satisfied. And the signal for a voxel *q* in an active region *i* is defined as ***y***_sig_(*q*) = ***X***_i_***β̃***_i_(*q*) where ***β̃***_i_(*q*) = ***β***_i_ + 0.1 ∗ rnd(3,1). In this equation, the term rnd(3,1) is a 3 x 1 vector generated from a standard normal distribution to introduce the variability between voxels. For the uniHRF simulation, the design matrix ***X***_i_ is the same across brain regions for a single subject since ***X***_i_ was obtained by convolving the binary task with the canonical HRF. In contrast, HRFs for the six active regions in each “subject” were randomly sampled from the set of HRFs plotted in Fig.4, where the white dashed line is the canonical HRF. We generated these HRFs with the *spm_hrf* function in SPM12 with all six parameters randomized: Delay of response (6+0.5*N(0,1)), delay of undershoot (16+N(0,1)), dispersion of response (1+0.2*N(0,1)), dispersion of undershoot (1+0.2*N(0,1)), ratio of response to undershoot (0+0.5*U(0,1)), and onset (0+0.3*U(0,1)), where N(0,1) is the normally distributed random number and U(0,1) is the uniformly distributed random number in the interval (0,1). The parameters without randomness generate the canonical HRF, the strength of randomness for each parameter is adjusted to have reasonable HRF shape. For both uniHRF and varHRF simulations, the resting-state time series were directly used for inactive regions. For active regions, the time series for each voxel was generated by adding resting-state time series ***y***_*rf*_ with a fraction of the signal, namely, ***y*** = ***y***_*rf*_ + *f* ∗ ***y***_sig_. The initial value of *f* was initialized with the value 0.2 and the GLM was applied to obtain the correlation map. Since the true activation status for each voxel is known, the false positive rate (FPR) and true positive rate (TPR) with varying correlation thresholds were calculated, and then receiver operating characteristic (ROC) curves were plotted to evaluate the overall sensitivity and specificity of generated fMRI data. The area under ROC curve (AUC) with FPR in the range from 0 to 0.1 (namely the maximal value of AUC is 0.1) was calculated. Since fMRI activation maps are usually thresholded with a strict p value to eliminate false positive voxels, the AUC value in the low FPR range 0-0.1 is more meaningful than the AUC value in the full FPR range 0-1 for controlling false positives. Consistent with our previous studies (Yang et al., 2018; Zhuang et al., 2017) having AUC value for fMRI data in the range of 0.05-0.07, the signal fraction *f* was adjusted if the simulated data with initial signal fraction value had an AUC value not in this range.

**Figure 4.**
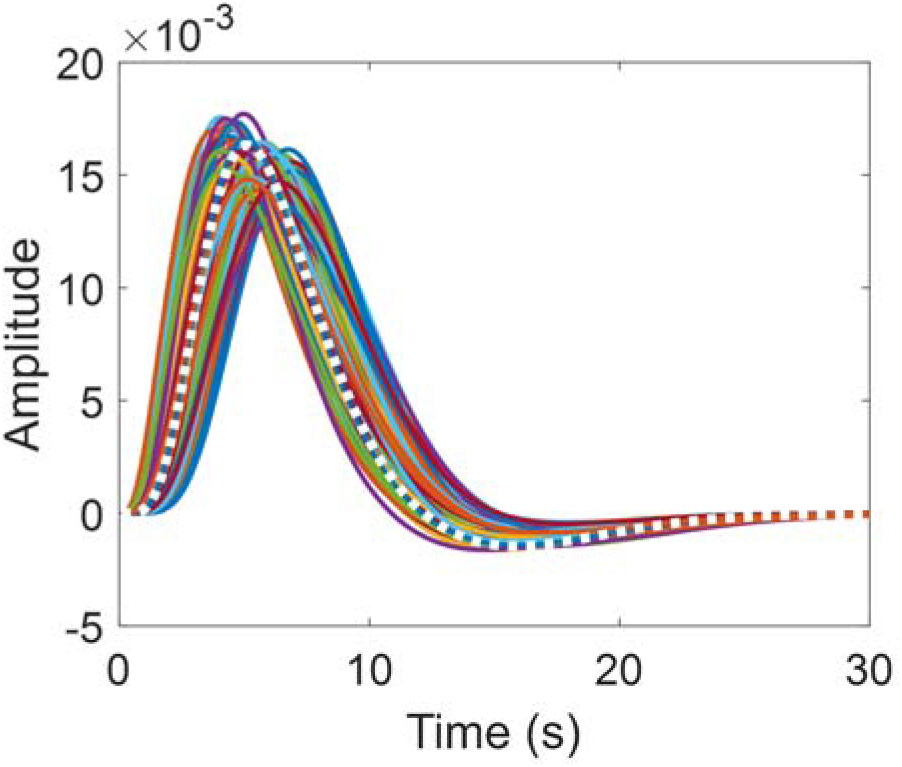
The set of hemodynamic response functions (HRF) used in the varHRF simulation. The white dashed line is the canonical HRF.

**Figure 5.**
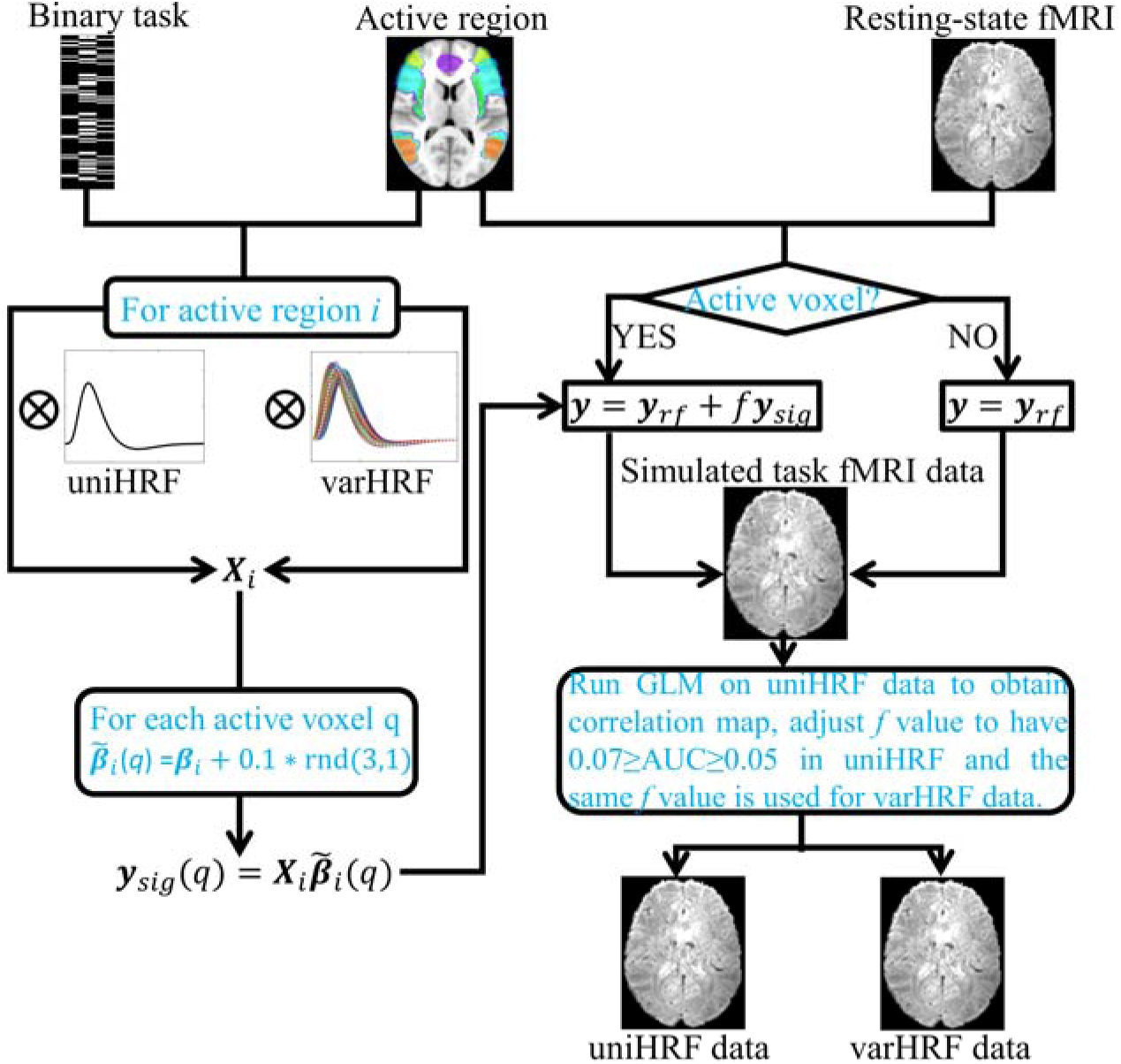
The flowchart to generate uniHRF and varHRF task fMRI data. The uniHRF data have the signal generated from a single canonical HRF, in contrast, varHRF has varying HRFs for different active regions. The rnd(3,1) is a 3 x 1 vector generated from standard normal distribution to introduce the variability between voxels within the same active region. The signal fraction *f* is initialized with the value 0.2 and adjusted to have uniHRF data having AUC within the range of 0.05 – 0.07. The varHRF data has the same signal fraction as uniHRF data for the same subject.

### 3.3. Data analysis

Following the tradition in fMRI activation analysis, the design matrix used in GLM was computed by convolving the binary task with the canonical HRF for both simulated and real fMRI data, regardless of the varying HRFs used to generate signal for the varHRF data. Multiple methods were used to process fMRI data, including the proposed *DNN* method, ICA-based denoising technique FIX with nuisance regression included and temporal filtering (Chai et al.). A detailed description about the proposed DNN method can be found in the *Theory* section and a brief description about FIX and TF is presented in the following.

The FIX tool (Griffanti et al., 2014) in FSL uses the output from ICA (MELODIC) to distinguish neural-related or noise-related components based on manually-labelled pre-trained data. The MELODIC ICA tool in FSL (v5.0.9) (see https://fsl.fmrib.ox.ac.uk/fsl/fslwiki/MELODIC) is commonly used along with other tools for the purpose of denoising fMRI data. MELODIC decomposes fMRI data into a set of independent components (ICs) with automatic dimensionality estimation and outputs filtered fMRI data. These ICs are then classified as either neural-related components or noise-related components. The classification can be either done automatically by machine learning methods or manually by hand labelling. A study by Griffanti et al. (2017) provides comprehensive guidelines for manual classification of ICs based on topological appearance of spatial maps, temporal power spectra, and certain features (such as spikes) in time courses. For HCP minimally-preprocessed 3T resting-state fMRI data, the MELODIC ICs were manually classified by experts and a pre-trained classifier is included in the FIX toolbox. Since the simulated task fMRI data were generated based on HCP resting-state data, it is reasonable to use the pre-trained classifier on the simulated data. The pre-trained classifier for HCP 3T resting-state data in FIX toolbox recently was shown also to be applicable for denoising HCP task fMRI data (Glasser et al., 2017), hence the pre-trained classifier is also used for denoising real working memory task fMRI data in our analysis. FIX was applied with multiple thresholds (e.g. 1, 5, 10, 20, 25, 30, 35, and 40) to separate neural-related ICs from nonneural (noise) ICs. A higher threshold intends to aggressively remove more non-neural components but may also remove neural-related components. For the simulated datasets, the threshold achieving maximal AUC value was chosen in the analysis. For the real data, we manually classified ICs from 10 subjects and observed that a threshold of 30 was a proper choice without any manually identified signal components classified as noise. The same threshold was applied for all the working memory task fMRI data.

Unless specifically mentioned, nuisance regression was applied along with FIX denoising by default. Nuisance regression is a prevalent preprocessing strategy to remove non-neural fluctuations in fMRI data. The nuisance regressors in this study are derived from estimated motion parameters during realignment of EPI data and their first-order derivative, and three principal components each from WM and CSF derived by using the CompCor algorithm (Behzadi et al., 2007; Muschelli et al., 2014), which efficiently captures physiological noise without external recordings. Band-pass temporal filtering (e.g. 0.01-0.1 Hz) is a commonly used technique for processing fMRI data. Considering that high-frequency fMRI signal (>0.1 Hz) recently gained attention in neuroimaging research community (Boubela et al., 2013) and a low frequency cutoff may partially remove meaningful neural signal. We have run analysis on uniHRF simulation data with high frequency threshold ranging from 0.10 Hz to 0.3 Hz with step 0.05Hz. The mean AUC values over 15 subjects was improved by 4% from 0.1 Hz to 0.15 Hz. Further increasing the frequency threshold up to 0.3 Hz has no considerable influence (less than 1%) on the result in terms of AUC values. Therefore, the frequency range 0.01-0.15 Hz was selected and used for our analysis.

For the DNN denoising method, the input fMRI data of size *N* x *T* x 1 are padded with zeros along the second dimension for the computations in the convolutional layer to make sure that the output has the same length of *T*. In our analysis, the convolutional layer is designed with 4 (*V*=4) 1-dim filters with a stride of 1 and filter size of 15. Thus, the first hidden layer has the dimension of *N* x *T* x 4. The LSTM layer has 4 units (*L*=4) and the hyperbolic tangent function is used as activation function. The number of nodes *K* in the time-distributed fully-connected layer varies in our study. The specific number of nodes are described in the result section for each case. *DNN* converges in 30 epochs for our simulated and real fMRI data. The computational time for each subject is less than 30 minutes on a Tesla K40c GPU with 2,880 CUDA cores. In the result section, five sets of different processed fMRI data for both simulated and real fMRI data were used for comparison, including raw (only minimally preprocessed and detrended) data and data processed by FIX, FIX+TF, *DNN* and FIX+*DNN*. The DNN denoising method was implemented with Theano deep learning package (http://deeplearning.net/software/theano/) and is publicly available (https://github.com/pipiyang/DeNN-task-fMRI-denoising).

## 4. RESULTS

### 4.1. Simulation

The uniHRF simulated data was analyzed with the DNN network using a single node (*K*=1) for the time-distributed fully-connected layer. For the varHRF simulated data, the DNN uses 4 nodes to adapt to varying HRFs among brain regions. Further increasing the LSTM nodes to 6 or 8 has negligible improvement for the varHRF simulation. Results for 5 different processed datasets are shown in Fig. 6 including just the raw data and data processed by FIX, FIX+TF, *DNN* and FIX+*DNN* for a simulated single subject. The time series for inactive voxels are shown in the first column and the voxel time series within 6 bilateral active regions are shown in the second to seventh columns. While the signals in each region *i* were generated with between-voxel variability, namely, ***y***_sig_(*q*) = ***X***_i_ ***β̃***_i_(*q*) where ***β̃*** _i_(*q*) = ***β***_i_ + 0.1 ∗ rnd(3,1), only the main signal ***y***_*sig*_ = ***X***_*i*_***β***_*i*_ is shown on the top panel of the figure. For the raw fMRI data, there exists prolonged noise fluctuation across the entire brain in the interval marked in blue. FIX efficiently removed the noticeable prolonged noise and the additional band-pass temporal filter TF improved the functional contrast-to-noise ratio (CNR) of the task effect in the data. Qualitatively, the *DNN* outperformed FIX and FIX+TF in improving the SNR, however, the prolonged noise fluctuations still exists in *DNN* processed data. The FIX+DNN processed data appeared to be the best denoised data, which have an improved CNR ratio and did not have any noticeable prolonged noise fluctuations.

**Figure 6.**
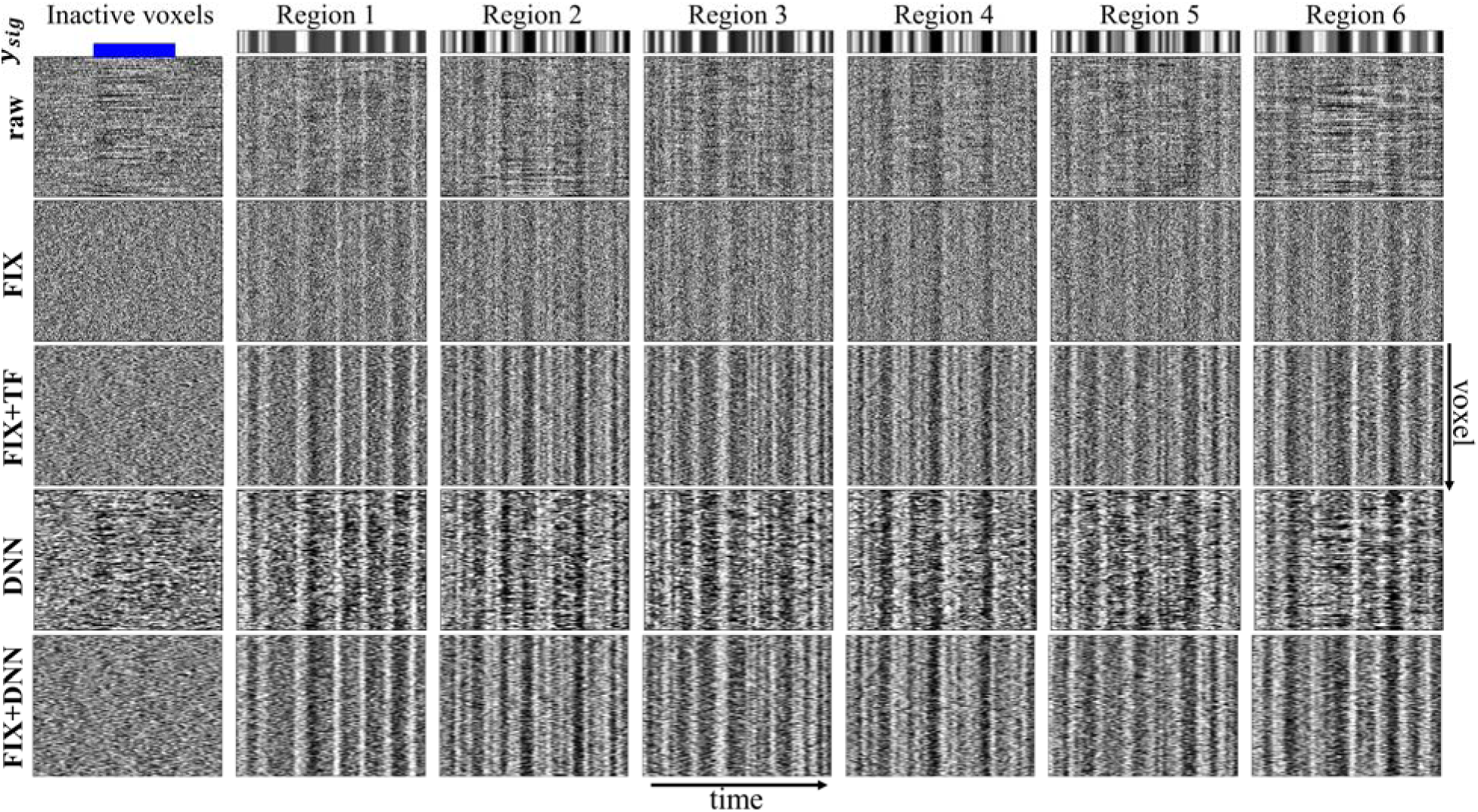
The raw, FIX, FIX+TF, *DNN*, FIX+*DNN* processed time series for a single “subject” chosen from the uniHRF simulation. The main signal ***y***_*sig*_ = ***X***_*i*_***β***_*i*_ is shown at the top panel for each region *i*. For the purpose of visualization, the height of each subplot is specified to be the same without reflecting the number of voxels in each region.

ROC curves were used to quantify the sensitivity and specificity in these five processed datasets. The FPR and corresponding TPR values for uniHRF and varHRF were calculated and the ROC curves are shown in Fig.7. The ROC curves show that the fMRI data with additional processing steps outperformed raw fMRI data in terms of activation detection. The percentage of AUC value %_AUC_within the FPR≤0.1 range defined as 

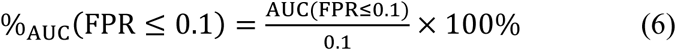

is used to quantitatively evaluate the performance of different techniques. For the uniHRF simulation, the %_AUC_ values are 62.4%, 67.8%, 72.4%, 69.2%, and 75.2% for raw, FIX, FIX+TF, *DNN*, FIX+*DNN* processed fMRI data. For the varHRF simulation, the %_AUC_values are 58.2%, 63.3%, 68.2%, 67.8% and 72.5% for processed data in the same order. In addition to ROC curves, the correlation difference between active and inactive voxels within gray matter mask were also calculated. The distributions of correlation for the five processed data in uniHRF simulation are shown in Fig.8, the distribution for inactive voxels are shown on the left and the distribution for active voxels are shown on the right. The varHRF simulation has similar distributions and are not shown in the figure. The mean correlation differences of raw data are 0.118 and 0.110 for uniHRF and varHRF, respectively. FIX+*DNN* has maximal correlation difference for both uniHRF and varHRF with values 0.259 and 0.244, respectively. The standard error of correlation differences is of the order of 10^-4^ and is not listed in the figure.

**Figure 7.**
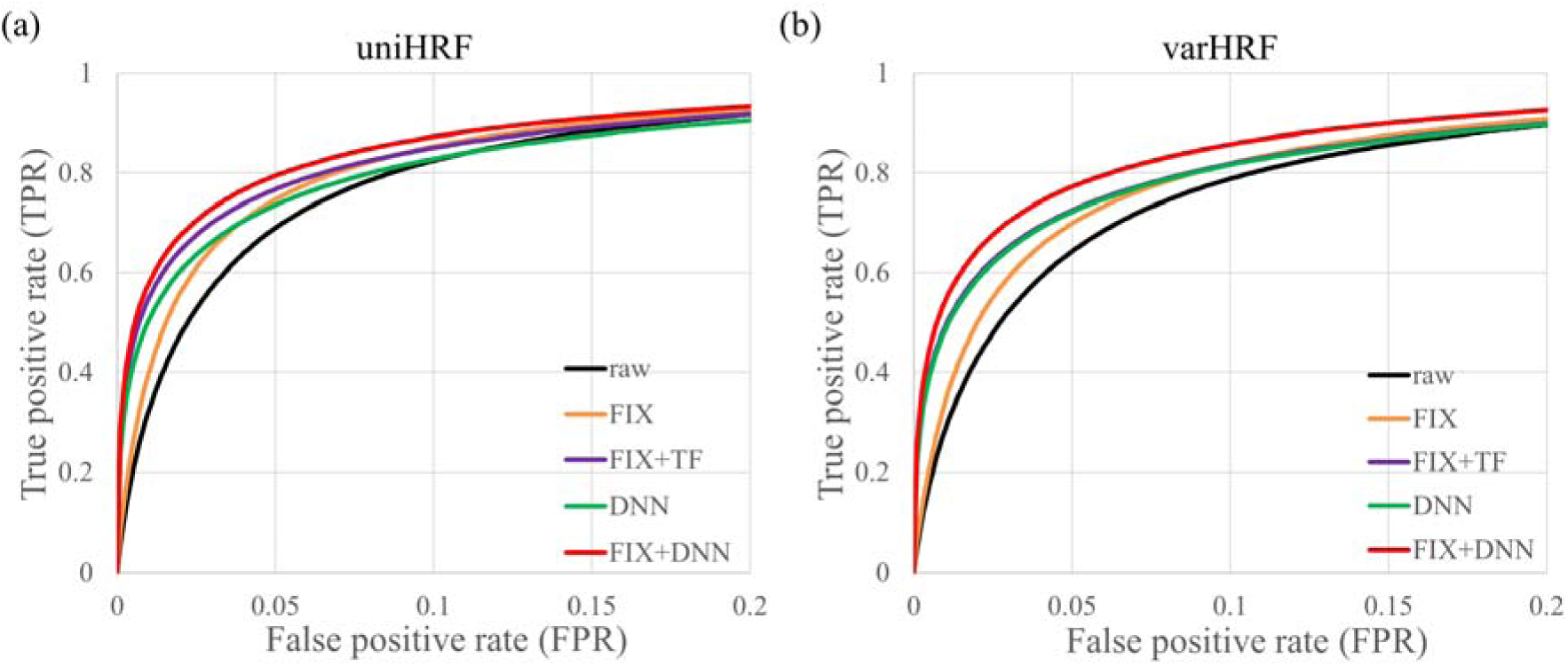
Receiver operating characteristic curves for raw, FIXneg, FIX+TF, *DNN* and FIX+*DNN* processed fMRI data. (a) uniHRF simulation, (b) varHRF simulation.

**Figure 8.**
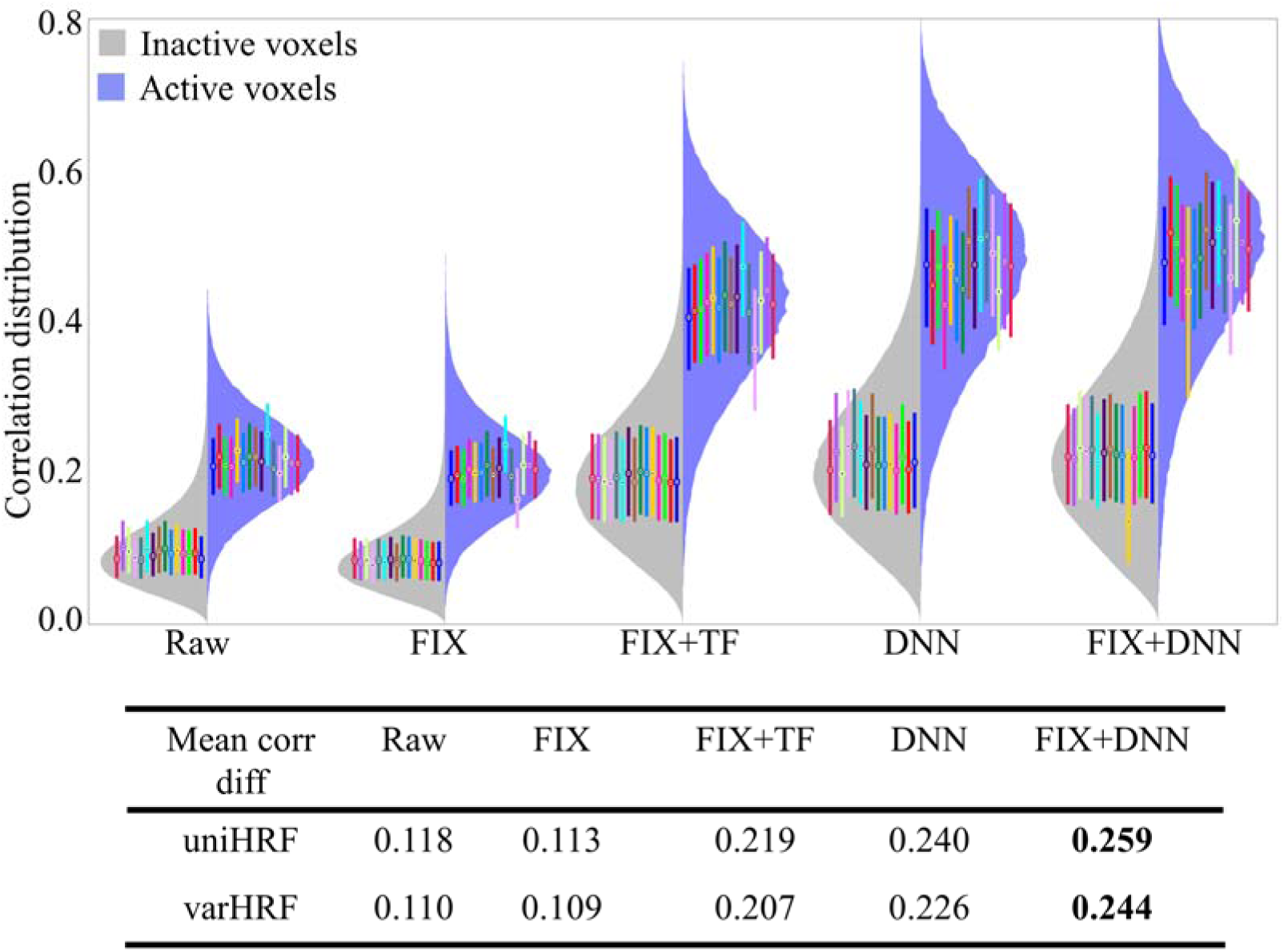
Distribution of correlations for different processed fMRI data for the uniHRF simulation. The distribution of inactive voxels over all 15 subjects is shown on the left side and the distribution of active voxels is shown on the right side. Not shown is the distribution for the varHRF simulation which is very similar to uniHRF. The compact boxplot of the inactive or active voxels for each subject is also shown in the figure with the bar indicating the range from 25^th^ percentile to 75^th^ percentile. Each color of the boxplot corresponds to a different subject. The mean correlation difference between inactive and active voxels for both uniHRF and varHRF simulations is listed in the figure. The standard error for all correlation differences is of the order of 10^-4^.

**Figure 9.**
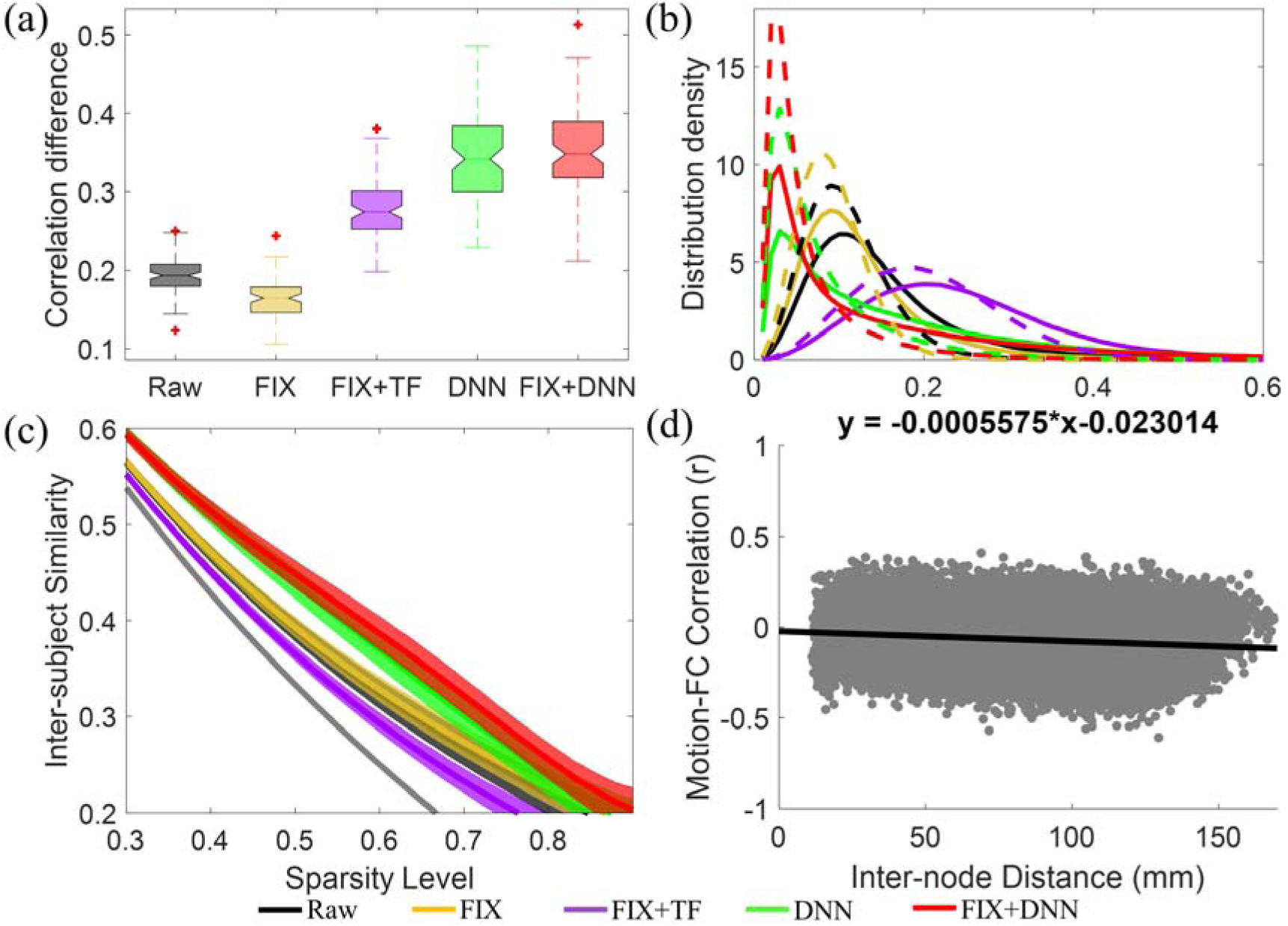
Results using HCP working memory fMRI data. (a) Boxplot of correlation difference between 90 percentiles and 10 percentiles of all correlations of voxel time series and task regressors in gray matter. (b) Distribution density of task-related correlation for gray matter (solid lines) and white matter voxels (dashed lines). (c) Similarity between subjects’ correlation maps at varying sparsity levels. (d) Correlation between pairwise connectivity (from raw data with 34,716 unique connections) and mean framewise displacement across subjects versus inter-node distance.

### 4.2. Real data

The methods applied on simulated data were also used to process real task fMRI data. Similar as in the uniHRF simulation, a single node was specified for the time-distributed fully-connected layer. Though there is no ground truth that defines activated voxels, our hypothesis is that voxels having lower average correlation with the design matrix are less likely to be active. The correlation difference between high-correlation and low-correlation voxels in real data can act as a criterion similar to the correlation difference between inactive and active voxels in the simulation. For each subject, the GM voxels were sorted in descending order in terms of the correlation value of each voxel time course with the design matrix. We have heuristically selected the 90 and 10 percentiles of the cumulative distribution of correlation values to calculate the correlation difference between active and inactive voxels for each subject. In Fig.9a we show a boxplot of this correlation difference for 88 subjects. The median values of the correlation differences were 0.194, 0.164, 0.275, 0.342, and 0.348 for raw, FIX, FIX+TF, DNN, and FIX+DNN processed data, respectively. Similar to simulated data, in real data FIX reduced the correlation difference, but FIX+TF, DNN and FIX+DNN increased the correlation difference. The distributions of correlations across subjects for all GM voxels (solid lines) and WM voxels (dashed lines) are shown in Fig.9b. The curves were normalized to have the area under the curve equal to one. GM voxels, as expected, overall have a stronger right tail for all processed datasets. Except for FIX+TF, all other methods pushed the peak of the distribution towards the negative direction (compared to the raw data).

Jaccard similarity coefficient is another measurement used to compute the similarity of correlation maps among all 88 subjects. Improved similarity between subjects’ activation maps is expected if a denoising technique correctly reduces the noise without considerably corrupting the signal in the data. In detail, the correlation maps were first binarized by thresholding the correlation value to achieve a sparsity (#of zero entries / #of total entries) varying from 0.1 to 0.9 in steps of 0.01. A higher sparsity means more zero elements. The Jaccard similarity coefficient (∈[0,1]) in our case is defined as the ratio of the number of non-zero entries in the intersections to the total number of non-zero entries. A coefficient closer to 1 means that the two maps are more similar. For each denoising method at each sparsity level, the Jaccard similarity is computed for each pair of subjects, leading to (88*87)/2=3,828 values for all subjects. In Fig.9c we show the median Jaccard similarity coefficient curves for raw (black), FIX (yellow), FIX+TF (purple), DNN (green) and FIX+DNN (Jorge et al.) and the 25^th^ and 75^th^ percentiles were used to generate the shaded area. We have also generated the null distribution of the Jaccard similarity coefficient by randomly permuting the binary entries for each subject. The median Jaccard similarity coefficient for the null maps is shown in gray line and the shared area was generated with p value of 10^-3^. The distribution is very narrow, thus the shaded area is barely visible in the figure. Compared to raw data, FIX slightly increased the Jaccard similarity coefficient and FIX+DNN overall had the largest coefficient. In contrast, FIX+TF reduced the Jaccard similarity coefficient.

We also investigated the effect of motion in our data. Since motion-related regressors were used in FIX, FIX+TF and FIX+DNN for processing fMRI data, the variance explained but these motion regressors cannot be used to measure motion contamination because otherwise motion assessment becomes a circular analysis which is invalid. Instead, distance-dependent functional connectivity is a commonly used approach to evaluate the influence of motion in resting-state fMRI data. We applied this technique on our task fMRI data. We used the atlas with 264 regions of interest (ROI) (Power et al., 2011), derived from the results of many task fMRI meta-analysis and a resting-state parcellation strategy (Cohen et al., 2008) to compute functional connectivity. The correlation between pairwise connectivity from raw data (34,716 unique connections) and mean framewise displacement (Power et al., 2012) across subjects were calculated. Similar to the distance-dependent correlation plot in Satterthwaite et al. (2013), the plot of the correlation versus inter-node Euclidean distance for raw data are shown in Fig.9d. Motion-FC correlation barely had association with inter-node distance even in the raw data. The other processed data had similar association plots, which were not shown in the figure.

Similar to the analysis performed by HCP site investigators (Barch et al., 2013), activation count maps (ACM) were generated to demonstrate the proportion of subjects that showed activation or deactivation at a z-threshold of 1.96 (uncorrected two-tailed p value 0.05) for the *targets* contrast [1 0 0]. The *target* contrast’s spatial map for each participant was binarized using this threshold and the ACM maps for each processed data were created by calculating the percentage of subjects having the common activation (red in Fig.10) and de-activation (blue in Fig.10) passing the threshold for each voxel. Overall, all ACM maps showed activation in regions including dorsal and ventral prefrontal cortex, dorsal parietal cortex, insula and anterior cingulate cortex and deactivation in regions including posterior cingulate cortex and Rolandic operculum. However, DNN and FIX+DNN processed datasets had more robust activation than the other three datasets in terms of cluster size and percentage value.

**Figure 10.**
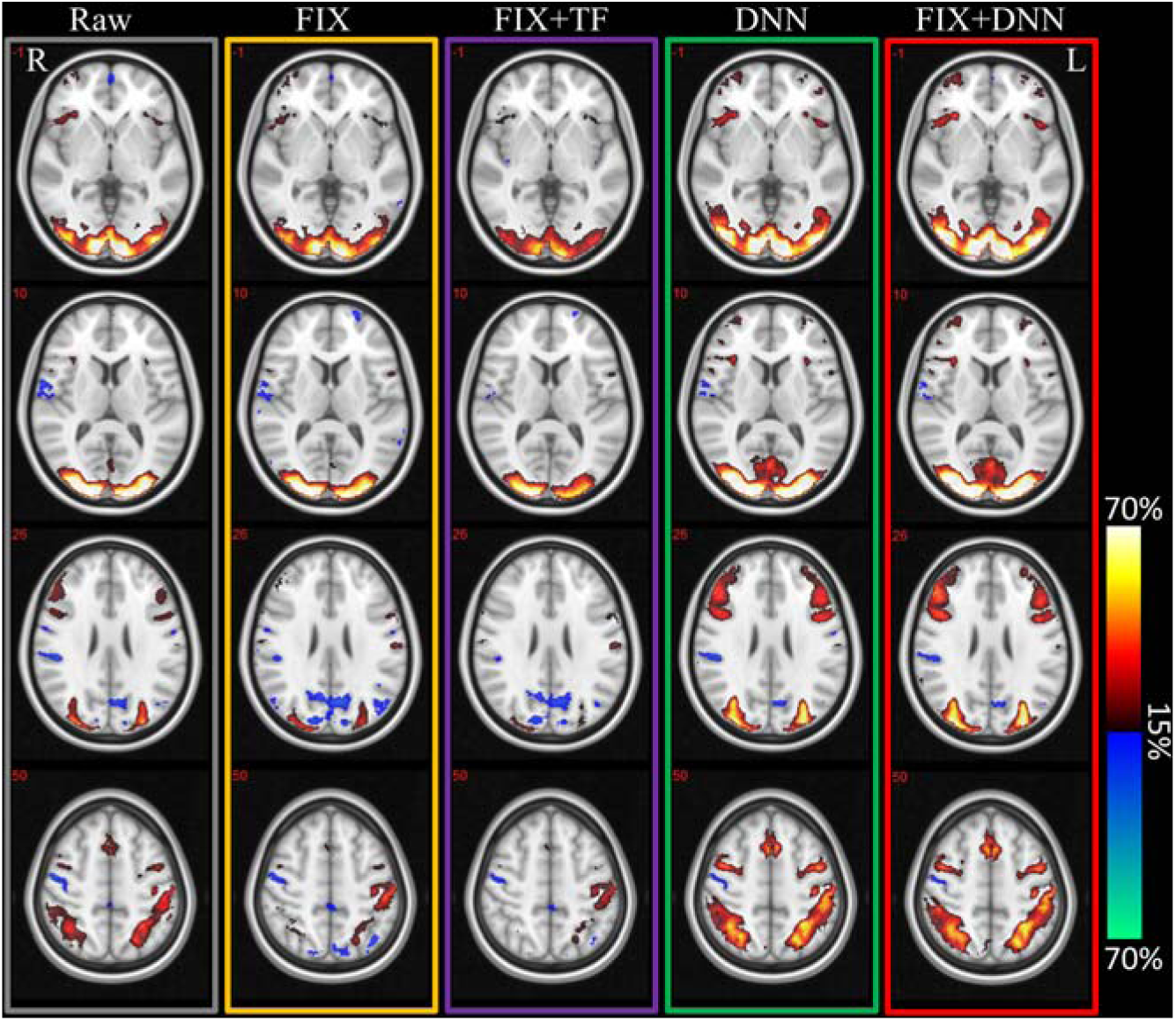
Activation count maps for raw, FIX, FIX+TF, DNN and FIX+DNN processed working memory task fMRI data for the *targets* contrast [1 0 0]. The targets contrast spatial map for each participant was binarized using a z-threshold of 1.96 (uncorrected two-tailed p value 0.05) and the ACM maps for each processed data were created by calculating the percentage of subjects having the activation (red regions) or de-activation (blue regions) passing the threshold for each voxel.

To visualize the influence of two intermediate layers, including convolutional layer and LSTM layer, on the frequency spectrum of time series, we have shown the estimated power spectral density (PSD) (Fulop and Fitz, 2006) of the output time series from these two layers from a single subject in Fig.11, in addition to the PSD of raw time series (the raw time series after 0.15 Hz low-pass filtering is shown in blue). The area under PSD curves are normalized to be 1. For the raw time series, the PSD for each voxel is first estimated and then the median PSD value at each frequency is used to generate the curve in the figure. The output time series from each filter in the convolutional layer are computed separately to generate the PSD vs frequency curve. The same process is applied to plot the PSD vs frequency plot for the LSTM layer. Naturally, there are *V*=4 curves for convolutional layer and *L*=4 curves for the LSTM layer. Compared to raw time series, the convolutional layer has reduced a large fraction of high frequency content in the raw time series. The filters in the convolutional layer, however, still show distinct frequency spectra, indicating large inter-filter variation. The LSTM layer further refines the frequency in the time series and achieves more similar PSD vs frequency curves across different LSTM units. Unlike low-pass filtering which only reduces spectral density beyond the frequency threshold, the DNN adjusts spectral density over the entire frequency range.

**Figure 11.**
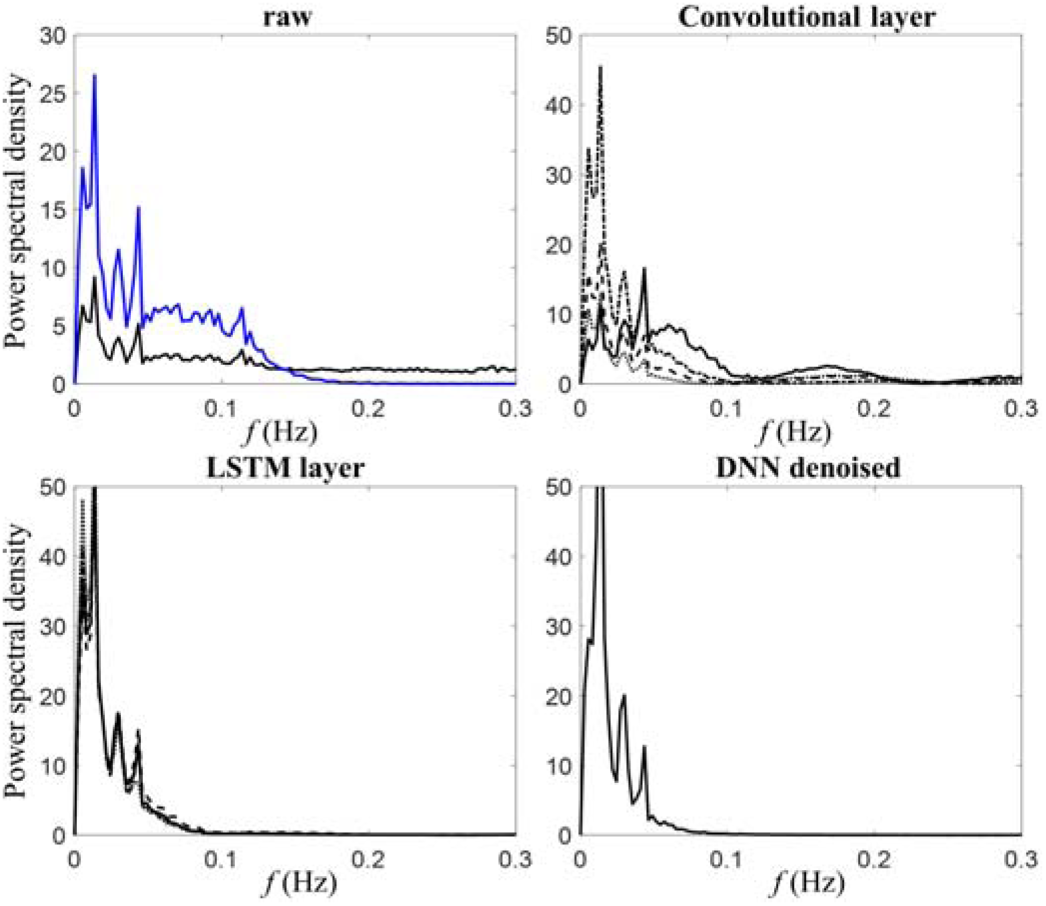
Estimated power spectral density for raw time series (for comparison a 0.15Hz low-pass filtered spectrum is shown in blue), and the output from convolutional layer and LSTM layer. The area under the curve is normalized to be 1. Note that the y-axis ranges are different for the raw time series.

Considering that LSTM units have similar frequency spectra, one question is whether we can replace the multiple LSTM units with a single LSTM unit. Since there is only a single output time series from LSTM layer for each voxel, the following time-distributed dense layer and selection layer are discarded and the output from the LSTM layer is treated as the denoised time series. We have run the analysis with the alternative setting (a single LSTM unit) on varHRF simulated data. The %_AUC_ is 64.7% for the alternative setting, in contrast, the proposed DNN framework has %_AUC_ as 67.8%. The distinct performance between these two settings indicates the necessity of the fully connected layer and selection layer. Furthermore, while LSTM units have similar PSD curves, having multiple LSTM units is still necessary to adequately describe arbitrary HRFs and variability of task response among voxels.

We used externally recorded physiological waveforms to evaluate how well the previously mentioned methods can reduce physiological noise Together with respiratory and cardiac regressors computed using the RETROICOR method (Glover et al., 2000; Kasper et al., 2017), respiration volume per unit time (RVT) regressors (shifted from -24 to 18 s with step=6 s) and cardiac-rate regressors (shifted from -12 to 12 s with step=6 s) (Bianciardi et al., 2009; Birn et al., 2008; Chang et al., 2009; Shmueli et al., 2007) were used to calculate the remaining physiological variance in fMRI data. The median variance explained by these physiological regressors were calculated for each subject. We show in Fig.12 a boxplot of the physiological variance in processed fMRI data. We separated the different processed fMRI datasets into a low-pass filtered (FIX+TF, DNN, FIX+DNN) and a *not low-pass* filtered (Raw and FIX) set. The raw fMRI data with 0.15 Hz low-pass filtering (Raw+TF) was also added to the low-pass filtered group as a reference for the other methods. With the low-pass filtered set, physiological noise accounted for a variance in the time series with median values of 18.4%, 13.6%, 3.4% and 2.2% for Raw+TF, FIX+TF, DNN and FIX+DNN, respectively. With the *not low-pass* filtered set, physiological noise accounted for a variance in the time series with median values of 3.7% and 0.3% for Raw and FIX, respectively. For both raw and FIX, the low-pass filtered datasets (Raw+TF and FIX+TF) had significantly higher physiological variance than their corresponding *no low-pass* filtered datasets, which indicates that physiological fluctuations have more variance in the low-frequency range. Among the low-pass filtered datasets, DNN and FIX+DNN had remaining physiological noise significantly (p < 10^-4^) lower than Raw+TF and FIX+TF.

**Figure 12.**
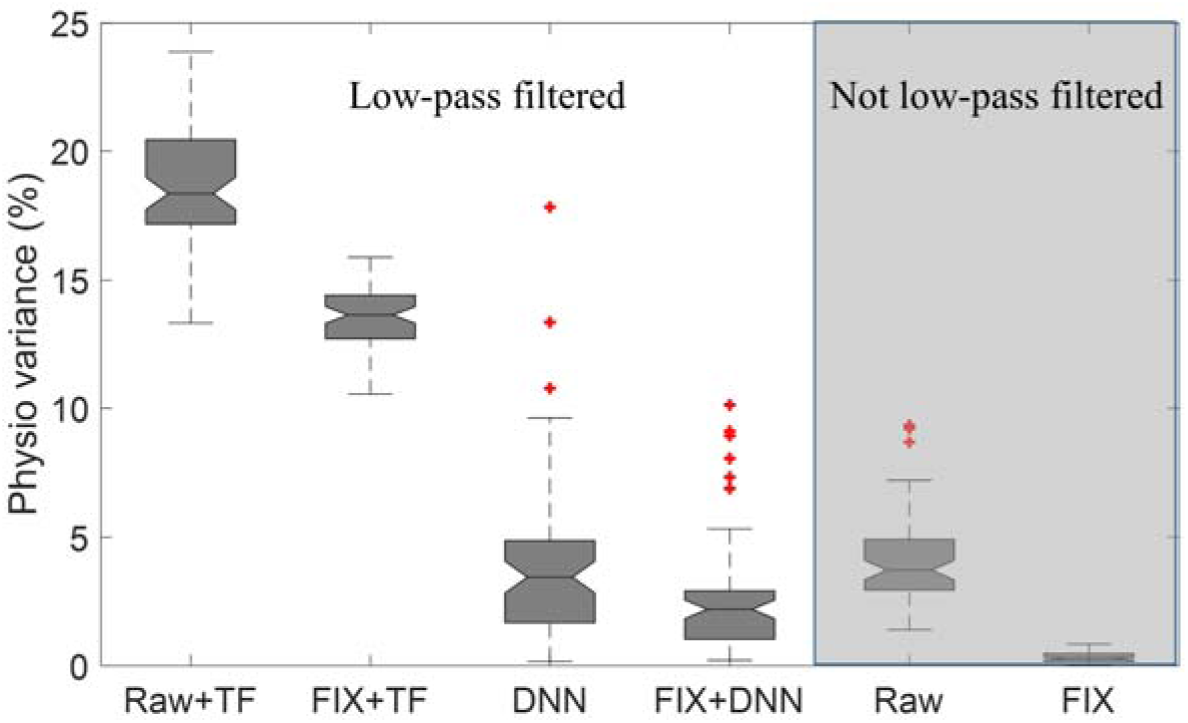
Physiological variance accounted for in different processed fMRI data. The raw fMRI data with low-pass filtering at 0.15 Hz (Raw+TF) was added to the low-pass filtered group to serve as a reference for the other methods.

## 5. DISCUSSION

In this study, we have designed a subject-level artificial neural network framework for denoising task-based fMRI data without assuming any explicit noise models. The proposed *DNN* network treats thousands of voxels as input samples to train the model separately for each subject. First, the raw fMRI data were temporally filtered by a 1-dim convolutional layer. The temporal filters in the convolutional layer are estimated from the data itself and do not correspond to conventional high-pass or low-pass frequency filters. Then, the LSTM layer was applied on the filtered time series from the convolutional layer to characterize the serial (auto) correlations of the temporal features in fMRI time series data. Finally, a time-distributed fully-connected layer combined with an unconventional *selection* layer is used to output the denoised time series. The *DNN* network has two input datasets, namely the time series within GM and nonGM voxels. These two input datasets share the same network (the same architecture with the same parameters) and the correlation difference between these two datasets with task-based regressors is maximized to optimize the parameters in the *DNN* network. Both simulated and real fMRI data suggest that DNN is a superior denoising technique and has the best performance when it was applied together with FIX denoising. To the best of our knowledge, this is the first study where a deep neural network is proposed for denoising task fMRI data.

### 5.1. Comparisons of different denoising methods

For the simulated data, while both DNN and band-pass filtering (Chai et al.) improved the CNR ratio, the DNN processed data achieved comparable performance with FIX+ band-pass filtering in activation detection. FIX (with nuisance regression included) complemented DNN in efficiently removing noise (see Fig.6). FIX+DNN outperformed the other single or dual denoising methods in uncovering task-related response. We quantitatively evaluated these methods by computing ROC curves and determining the correlation difference between inactive and active voxels. Compared to raw data, FIX+DNN improved AUC value by 20.5% and 24.4% for the simulations with signals generated with a single or multiple HRFs, namely uniHRF and varHRF, respectively. Even though varHRF and uniHRF were generated at the same signal to noise fraction, varHRF had lower AUC values because of the varying HRFs in different brain regions, regardless which method was used for denoising. The varHRF data processed without DNN had a lower AUC by approximately 4%, in contrast, the data processed with DNN (including DNN and FIX+DNN) had a lower AUC by only about 2%. We attribute the better performance of the DNN to the temporal convolutional layer and the LSTM layer. The convolutional layer automatically creates filters that can adapt to HRFs with different delays or onsets, longer or shorter plateaus and varying undershoot across the brain, and the LSTM denoises the current time point with information from previous time points. The robustness in performance of the DNN despite heterogeneous HRFs reduced potential misspecification of the modeled design matrix. Given the well-known occurrence of varying HRFs in brain regions, this adaptive property of the DNN is particularly useful for denoising.

We have used the correlation difference to evaluate how well different methods distinguish active voxels from inactive voxels. Both TF and DNN methods increased the correlation for all voxels including active and inactive ones in FIX+TF, DNN and FIX+DNN processed data. Furthermore, the AUC value improved for FIX+TF, DNN and FIX+DNN. In contrast, the FIX method without TF or DNN slightly decreased the correlation difference, but nonetheless, it improved the AUC value. We attribute increased AUC value to the reduced standard deviation of correlation values. Because ICA decomposition does not ensure that the time courses of ICA components are orthogonal (Caballero-Gaudes and Reynolds, 2017), regressing out these ICs may partially remove neural signal, which leads to decreased correlation difference. Choosing a more conservative threshold in FIX can reduce the risk of removing signal components with the drawback of removing less noise in the data. In addition, nuisance regression can also partially remove task-related fluctuations because of partial frequency overlap of signal and noise sources.

Similar to the findings in the simulation, an increased correlation difference in FIX+TF, DNN, FIX+DNN and decreased value in FIX was observed for the working memory fMRI data. With the hypothesis that denoising could make activation maps more homogeneous, we expected more similar correlation maps between subjects and used Jaccard similarity coefficients to measure the similarity for these processed datasets at varying sparsity levels. DNN had significant larger between-subject similarity coefficient and FIX+DNN overall achieved the largest similarity coefficient, indicating that the denoised data become more homogeneous with DNN. Applying FIX alone improved the between-subject similarity, but the additional band-pass filtering significantly lowered the similarity coefficient, which may be related to the frequency threshold and aliasing of different physiological noise sources. As a non-specific data-driven denoising method, the proposed DNN method did not target a particular noise source. By calculating how much physiological variance is left in the denoised data, we found that the proposed DNN method could significantly reduce physiological noise. We also investigated to measure DNN’s influence on motion effect, the raw data showed a negligible trend between motion-FC correlation and inter-node distance (see Fig.9d) and hence no further test was applied. The majority of the data had a mean framewise displacement less than 0.25mm which does not necessitate motion correction. In addition, task fMRI data has stronger neural response and higher signal-to-noise ratio than resting-state data. The task-related neural response may dominate over motion-related fluctuations. These two factors could explain why there is no significant trend in the plot of motion-FC correlation versus inter-node distance. However, we showed that DNN is also applicable and effective to reduce noise on more severe motion-corrupted fMRI data in Appendix A.

### 5.2. Characteristics of DNN denoising

While the denoising methods including ICA-based denoising (e.g. FIX), nuisance regression, temporal band-pass filtering (Chai et al.) and the proposed DNN aimed at improving fMRI data quality, these methods have different properties, assumptions and limitations. Before discussing the characteristics of the DNN method in detail, we would like to briefly summarize the other methods.

ICA-based denoising provides an efficient way to clean up structured noise components that may exhibit similar temporal characteristics but distinct spatial patterns compared to real neural activity, e.g. aliased physiological noise, motion effects, scanner artifacts and other nuisance resources. Nuisance regression was applied following FIX denoising with regressors for rigid-body head movement realignment parameters and their first order derivatives, physiological noise via CompCor, and the average signals of white matter and ventricular cerebrospinal fluid tissues. However, rigid-body movement estimation only approximately determines motion signatures, and thus nuisance regression cannot properly address motion-related signal changes that are not linearly related to the estimated realignment parameters and their derivatives or squares. Furthermore, the selection of nuisance regressors is another factor influencing the performance of fMRI denoising. The inclusion of global signal regression has been heavily debated in the recent past (Murphy et al., 2009; Power et al., 2018; Saad et al., 2012; Weissenbacher et al., 2009). Temporal band-pass filtering keeps a certain frequency band and discards the very low (drift frequency range) and high frequencies assumed to be irrelevant to the task. Naturally, the filtering cannot separate noise from the signal within the frequency band. In addition, the proper frequency band which depends on the form of the proper HRF is still under debate (Boubela et al., 2013).

The proposed DNN network takes advantage of the temporal correlations between adjacent time points to better extract the task-related neural response in fMRI with the assumption that no task-related neural activity exists within white matter and cerebrospinal fluid tissues. Compared to band-pass filtering, DNN can be treated as an adaptive filter that is determined by the data. This property makes DNN not only reduce high-frequency noise fluctuation in the data but also potentially separate the noise within the frequency range of the task-related neuronal signal. While both FIX and DNN involve training an algorithm or network for denoising, they have their own unique properties. If a pre-trained classifier is not available, FIX first trains the algorithm using hand-labelled signal and noise components from a few subjects and then applies the algorithm to classify noise components on new subjects. In contrary, DNN does not require hand labelling and is trained to have a unique model (the same framework but unique model parameters) for each subject. However, the DNN network does not consider structural noise information and thus it has difficulty in removing structured noise that exhibit similar temporal characteristics but distinct spatial patterns with the task-related neural activity, such as aliased physiological noise and task-introduced movement artifacts. This limitation is consistent with our result that the combination of FIX and DNN achieved best performance. Certainly, ICA-AROMA (Pruim et al., 2015) could be a replacement of FIX if motion-induced structural noise is the major concern. DNN uses the task design for denoising and is robust to HRFs varying with brain regions. The GLMdenoise proposed in Kay et al. (2013) also uses a task design for denoising and estimates subject-specific HRFs for analysis. GLMdenoise, however, requires multiple runs for the same task on a single subject, which is costly and usually not available in most studies. Since a task design is used to guide the denoising process, both DNN and GLMdenoise cannot differentiate noise from the signal that is not modeled in the task design. Therefore, if a researcher is interested in another set of task design for the same fMRI datasets, it is advisable to re-run DNN or GLMdenoise for denoising. Nuisance regression-based denoising methods are known to reduce the degrees of freedom (dof) in fMRI data and the statistical analysis needs to be adjusted to reflect this change. In contrast, the DNN denoising method does not change the dof. The number of dof equals the number of principal components that can be extracted by applying principal component analysis on whole brain time series. The number of principal components after DNN denoising was found to be the same as the original data.

In addition to the HCP data (acquired with a multiband EPI sequence, 405 volumes, TR=0.72s), we have also applied DNN denoising on block-design episodic memory task fMRI data acquired with a standard EPI sequence (288 volumes, TR=2s). The result (see Appendix A) showed that DNN processed data showed stronger activation in hippocampus and nearby regions, indicating that DNN denoising is robust for different task designs and EPI sequences, regardless of the considerable motion-related artifacts in the raw time series.

Setting the hyperparameters such as number of hidden layers and units, learning rate, batch size, and the number of epochs is an important topic in the deep learning community. The commonly used grid search technique in traditional machine learning becomes too time-consuming for deep neural networks because of the increasing number of hyperparameters. The hyperparameters used in this study are heuristically specified. Our experience suggests that the learning rate, batch size, and the number of epochs used in this study are quite generalizable for different data and specifying the filter size for the 1-dim convolutional layer as an integer close to 10/TR is a proper choice. The episodic memory task fMRI data discussed in previous paragraph was denoised using the same architecture as the HCP data, except a smaller filter size 10/TR=5 was used.

Resting-state data have gained tremendous attention in the last decade, and one interesting question is whether the DNN denoising can also be applied to resting-state data. Unfortunately, the current DNN framework cannot be directly applied to resting-state fMRI data, because there is no task design matrix and it is unclear what matrix could be the replacement in the resting-state. We have attempted to replace the task design with motion regressors, however, motion confounds fMRI data across the entire brain and thus the loss function to maximize the correlation difference between gray matter and non-gray matter is not valid.

## 6. CONCLUSION

We have proposed a deep neural network framework for denoising task-based fMRI data and compared the *DNN* network with other denoising methods using simulated and real fMRI data. The *DNN* network significantly increased the correlation difference between inactive and active voxels and improved the activation detection in the simulation. Results from real data showed that DNN can efficiently reduce physiological fluctuation resulting in data that are more homogeneous across subjects. To the best of our knowledge, this is the first study using a deep learning algorithm for denoising task fMRI data.

## 7. ACKNOWLEDGEMENTS

This research project was supported by the NIH (grant 1R01EB014284 and COBRE grant 5P20GM109025), a private grant from Angela and Peter Dal Pezzo, and a private grant from Lynn and William Weidner. Data collection and sharing for this project was provided by the Human Connectome Project (HCP; Principal Investigators: Bruce Rosen, M.D., Ph.D., Arthur W. Toga, Ph.D., Van J. Weeden, MD). HCP funding was provided by the National Institute of Dental and Craniofacial Research (NIDCR), the National Institute of Mental Health (NIMH), and the National Institute of Neurological Disorders and Stroke (NINDS). HCP data are disseminated by the Laboratory of Neuro Imaging at the University of Southern California.

## Appendix A

Considering that the HCP data is not severely affected by motion, we have tested DNN denoising on motion-corrupted episodic memory task fMRI data acquired from seven healthy elderly subjects in another study.

FMRI data of seven elderly subjects were acquired with Institutional Review Board approval from University of Colorado, Boulder, on a 3T GE HDx MRI scanner equipped with an 8-channel head coil. Acquisition parameters for the EPI sequence were: TR/TE=2000 ms/30 ms, parallel imaging factor=2, slices=25 (coronal oblique, perpendicular to the long axis of hippocampus), slice thickness/gap = 4.0 mm/1.0 mm, 288 time frames (total scan duration 9.6 min), in plane matrix 96 x 96 voxels, FOV=220 mm. The fMRI volumes were interpolated to obtain an isotropic voxel size of 2 mm x 2 mm x 2 mm. A conventional structural T1-weighted image (0.43 mm x 0.43 mm x 1 mm) and a standard T2-weighted image (coplanar to the EPI) but with higher resolution (0.43 mm x 0.43 mm x 2.5 mm) were also acquired.

An episodic memory task was performed to obtain fMRI data. The episodic memory task contained visual stimuli, which show a novel face paired with an occupation. The entire task consisted of six periods of encoding, distraction, recognition and brief instructions to remind subjects of the task ahead. Specifically, during the encoding task, the subject was asked to memorize 7 faces paired with occupations, displayed in sequential order for a duration of 3s each and 21s in total. A distraction task (duration 11s) then followed each encoding task, where the subject was instructed to press the right or left button as fast as possible when the letter “Y” or “N” randomly appeared on the screen (right button for “Y” and left button for “N”). The recognition task consisted of fourteen stimuli, half novel and half identical to the stimuli seen in the previous encoding task. The subject was instructed to press the right button when the stimulus was previously shown and the left button when the stimulus was new. Scan duration was 9 min 36 s, and 288 time frames were collected. The task-related regressor ***X*** was constructed by convolving the task design consisting of 4 regressors for Instruction, Distraction, Encoding, and Recognition with the canonical hemodynamic response function.

The top panel in Fig.A1 shows the frame-wise displacement (FD) (Power et al., 2012) derived from six motion parameters for the seven subjects. The raw and DNN-denoised time series are presented on the second and third panel, respectively. The task design is plotted on the bottom panel. The raw fMRI data was observed to have considerable motion-related artifacts, in contrast, the DNN-denoised time series show more task-related temporal dynamics. We have applied GLM on raw and DNN-denoised time series to calculate the statistical activation map for the contrast “encoding-distraction” ([0, -1, 1, 0]). The activation maps at significance level *p*<10^-4^ in the standard MNI space are shown in Fig.A2a. The DNN has correctly detected the activation at the bilateral hippocampus and fusiform gyrus. We have also calculated the median correlations for the 50 percentile of voxels with high correlation and the other 50 percentiles with low correlation. With the hypothesis that the voxels with high correlation have more signal than the ones with low correlation, a good denoising method should boost the correlation for the top 50 percentiles and have less impact on the bottom 50 percentiles of voxels. The correlation difference obtained from raw and denoised fMRI data for the seven normal subjects are shown in Fig.A2b. Across all seven subjects, DNN has a much higher correlation difference than raw fMRI data.

**Figure A1.**
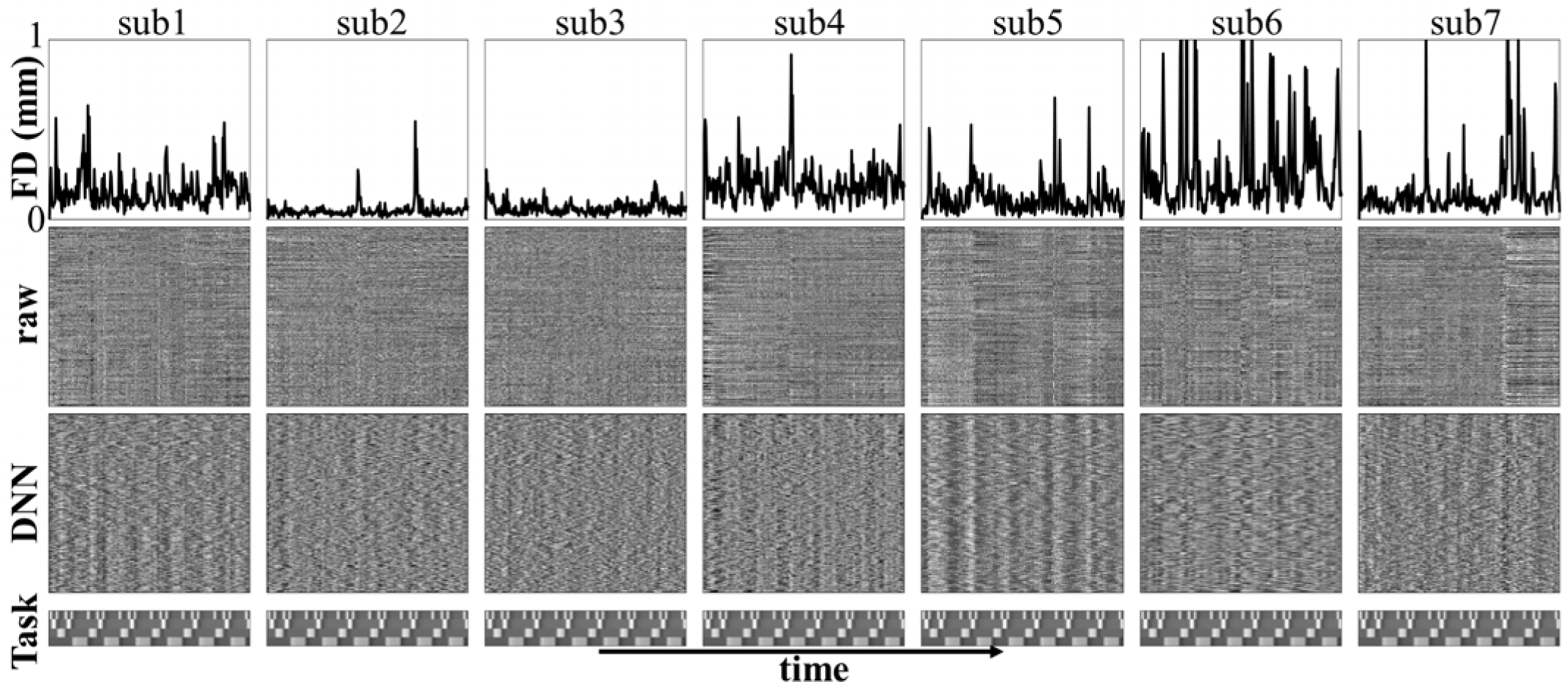
DNN denoising on motion-corrupted episodic memory task fMRI data from seven healthy elderly subjects. Framewise displacement is shown in the top panel. The raw and DNN denoised time series are in the second and third panel, respectively. The bottom panel shows the task design including instruction, distraction, encoding and recognition.

**Figure A2.**
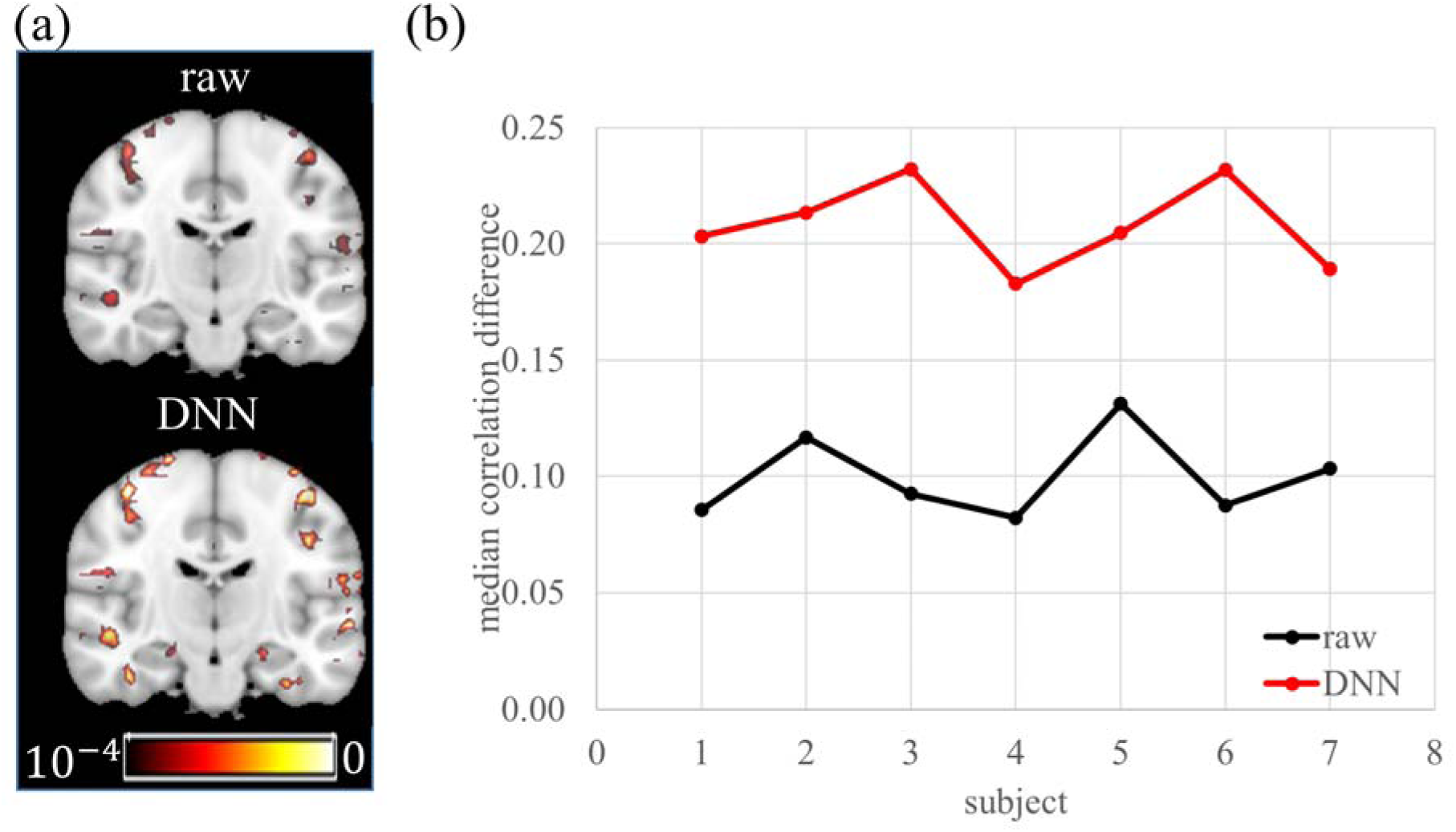
(a) Statistical activation map with the contrast “encoding-distraction” ([0, -1, 1, 0]) at significance level *p*<10^-4^. (b) Median correlation differences between the 50 percentiles of voxels with high correlation and the other 50 percentiles with low correlation for seven normal subjects.

**Zhengshi Yang** received the B.S. degree in Physics from Huazhong University of Science and Technology, Wuhan, China, in 2013 and the M.S. degree in Physics from University of Kentucky, Lexington, United States in 2015. Currently, he is a research engineer at the Cleveland Clinic Lou Ruvo Center for Brain Health in Las Vegas, United States. His research interests include multivariate mathematical modeling and MRI image analysis.
**Xiaowei Zhuang** received the B.S. degree in Biomedical Engineering from Jiaotong University, Beijing, China, in 2012 and the M.S. degree in Biomedical Engineering from University of Southern California, Los Angeles, United States, in 2014. Currently, she is a research engineer at the Cleveland Clinic Lou Ruvo Center for Brain Health in Las Vegas, United States. Her research interests include MRI image analysis and machine learning.
**Karthik Ramakrishnan Sreenivasan** received the M.S. degree in Electrical Engineering from Auburn University, Alabama, USA. He is currently a research engineer at the Cleveland Clinic Lou Ruvo Center for Brain Health in Las Vegas, Nevada, USA. His research interests include signal and image processing, investigation of brain connectivity using Functional Magnetic Resonance Imaging (fMRI) data, analysis of Positron Emission Tomography (PET) imaging data, and using machine learning approaches for brain state classification.
**Virendra Radheshyam Mishra** received the Ph.D. degree in Biomedical Engineering from joint universities of The University of Texas at Southwestern Medical Center at Dallas and The University of Texas at Arlington, Texas, USA and M. S. degree in Electrical Engineering from The University of Texas at Arlington, Texas, USA. He is currently a staff member at the Cleveland Clinic Lou Ruvo Center for Brain Health in Las Vegas, Nevada, USA. His research interests include image and signal processing of diffusion MRI and pseudo-continuous arterial spin labeling MRI data and predicting imaging biomarkers using machine learning techniques in clinical populations.
**Tim Curran** received the B.S. degree in Psychology from the University of Wisconsin-Madison in 1988, and M.S. and Ph.D. degrees in Psychology from the University of Oregon in 1990 and 1993, respectively. He then joined Harvard University for a postdoctoral fellowship in cognitive neuroscience from 1993-1995. His research interests include learning, memory, and visual object recognition. He is currently Professor of Psychology and Neuroscience at the University of Colorado Boulder.
**Dietmar Cordes** received the B.S. and M.S. degree in Physics from the Technical University of Clausthal, Germany, in 1981 and 1985, respectively, and in 1989 a Ph.D. degree in Theoretical Physics from the University of Nevada, Reno. He then joined the University of Wisconsin-Madison for a 4-year program in medical physics with focus on MRI and functional MRI (fMRI). His research interests include MR physics, MR imaging and data analysis in fMRI. He is currently the Director of Neuroimaging at the Cleveland Clinic Lou Ruvo Center for Brain Health in Las Vegas, NV, USA.

